## Supplementary Information for "Population genomics reveals an ancient origin of heartworms in canids"

Supplementary Fig. 1: Raw nuclear sequencing coverage per sample (Part 1/2). 3
Supplementary Fig. 2: Raw nuclear sequencing coverage per sample (Part 2/2). 4
Supplementary Fig. 3: Sex determination of samples based on relative coverage of sex-linked vs autosomal chromosomes (Part 1/2). 5
Supplementary Fig. 4: Sex determination of samples based on relative coverage of sex-linked vs autosomal chromosomes (Part 2/2). 7
Supplementary Fig. 5: Principal component analysis (PCA) of 196,550 nuclear single nucleotide polymorphisms (SNPs) from 124 heartworms with labelled metadata. 9
Supplementary Fig. 6: Principal component analysis (PCA) of 124 heartworms from around the world based on the nuclear genome 10
Supplementary Fig. 7: Genetic relatedness of the mitochondria and Wolbachia endosymbionts of heartworms 12
Supplementary Fig. 8: Admixture analysis of adult heartworms. 14
Supplementary Fig. 9: Population genetic summary statistics of heartworms collected from diverse host populations 15
Supplementary Fig. 10: Population genetic summary statistics of heartworms between broad geographical regions. 16
Supplementary Table 1: Adult heartworm burdens in Carnivora hosts 17
Supplementary Fig. 11: TreeMix analysis of nuclear heartworm data 22
Supplementary Fig. 12: Summary diagram of the population genomics workflow in this study 23
Supplementary Fig. 13: Demographic history analysis of heartworms from dogs using a range of generation times. 24

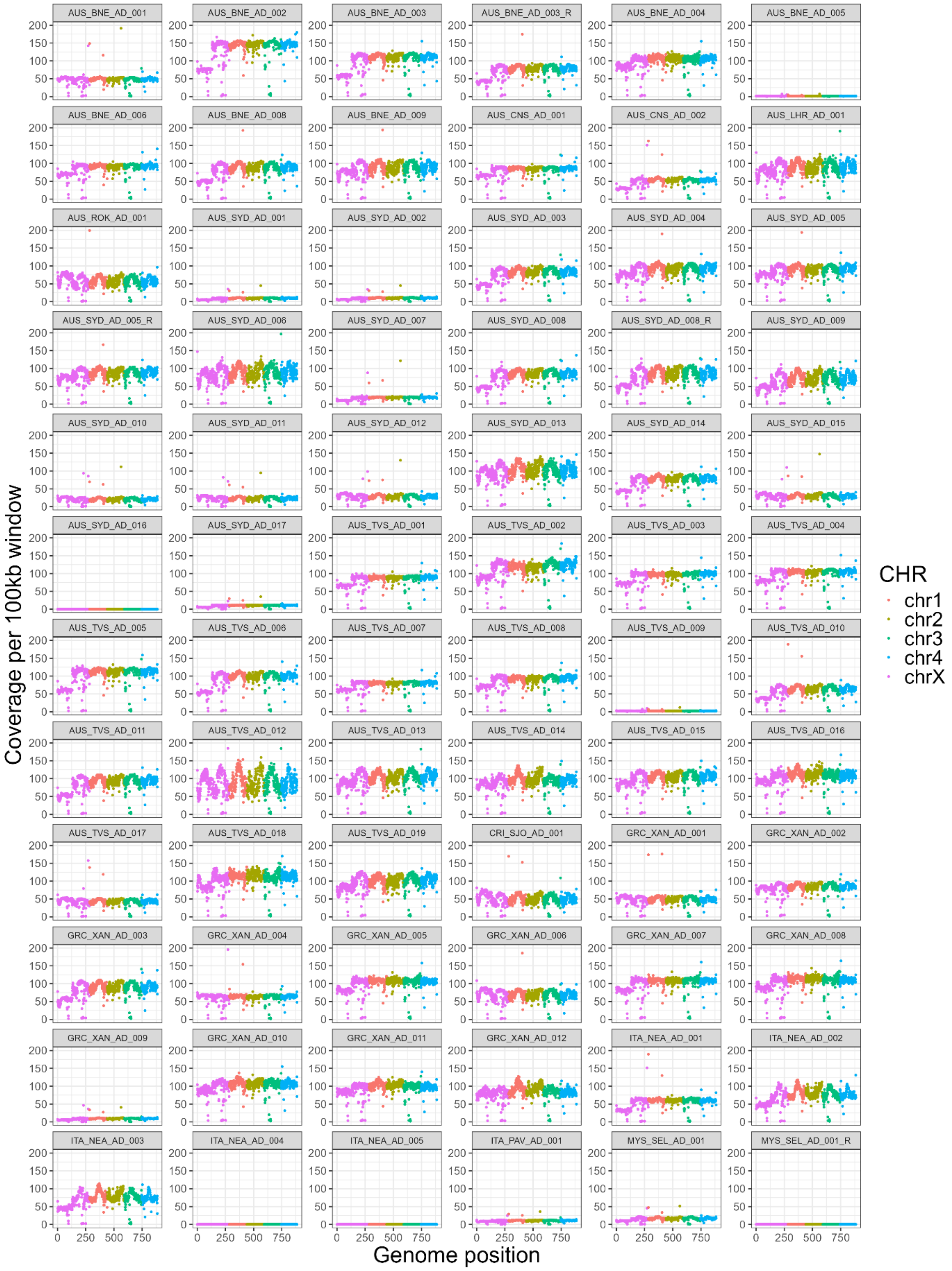
AUS_BNE_AD_001
AUS_BNE_AD_002
AUS_BNE_AD_003
AUS_BNE_AD_003_R
AUS_BNE_AD_004
AUS_BNE_AD_005
AUS_BNE_AD_006
AUS_BNE_AD_008
AUS_BNE_AD_009
AUS_CNS_AD_001
AUS_CNS_AD_002
AUS_LHR_AD_001
AUS_ROK_AD_001
AUS_SYD_AD_001
AUS_SYD_AD_002
AUS_SYD_AD_003
AUS_SYD_AD_004
AUS_SYD_AD_005
AUS_SYD_AD_005_R
AUS_SYD_AD_006
AUS_SYD_AD_007
AUS_SYD_AD_008
AUS_SYD_AD_008_R
AUS_SYD_AD_009
AUS_SYD_AD_010
AUS_SYD_AD_011
AUS_SYD_AD_012
AUS_SYD_AD_013
AUS_SYD_AD_014
AUS_SYD_AD_015
AUS_SYD_AD_016
AUS_SYD_AD_017
AUS_TVS_AD_001
AUS_TVS_AD_002
AUS_TVS_AD_003
AUS_TVS_AD_004
AUS_TVS_AD_005
AUS_TVS_AD_006
AUS_TVS_AD_007
AUS_TVS_AD_008
AUS_TVS_AD_009
AUS_TVS_AD_010
AUS_TVS_AD_011
AUS_TVS_AD_012
AUS_TVS_AD_013
AUS_TVS_AD_014
AUS_TVS_AD_015
AUS_TVS_AD_016
AUS_TVS_AD_017
AUS_TVS_AD_018
AUS_TVS_AD_019
CRI_SJO_AD_001
GRC_XAN_AD_001
GRC_XAN_AD_002
GRC_XAN_AD_003
GRC_XAN_AD_004
GRC_XAN_AD_005
GRC_XAN_AD_006
GRC_XAN_AD_007
GRC_XAN_AD_008
GRC_XAN_AD_009
GRC_XAN_AD_010
GRC_XAN_AD_011
GRC_XAN_AD_012
ITA_NEA_AD_001
ITA_NEA_AD_002
ITA_NEA_AD_003
ITA_NEA_AD_004
ITA_NEA_AD_005
ITA_PAV_AD_001
MYS_SEL_AD_001
MYS_SEL_AD_001_R
Coverage per 100kb window
0
50
100
150
200
Genome position
0
250
500
750
CHR
chr1
chr2
chr3
chr4
chrX

**Supplementary Fig. 1: Raw nuclear sequencing coverage per sample (Part 1/2).**

Coverage was calculated using 100 kb sliding windows for all mapped samples. Each chromosome is shown in a different colour, with a legend on the right. The remaining samples are shown in Supplementary Fig. 2.

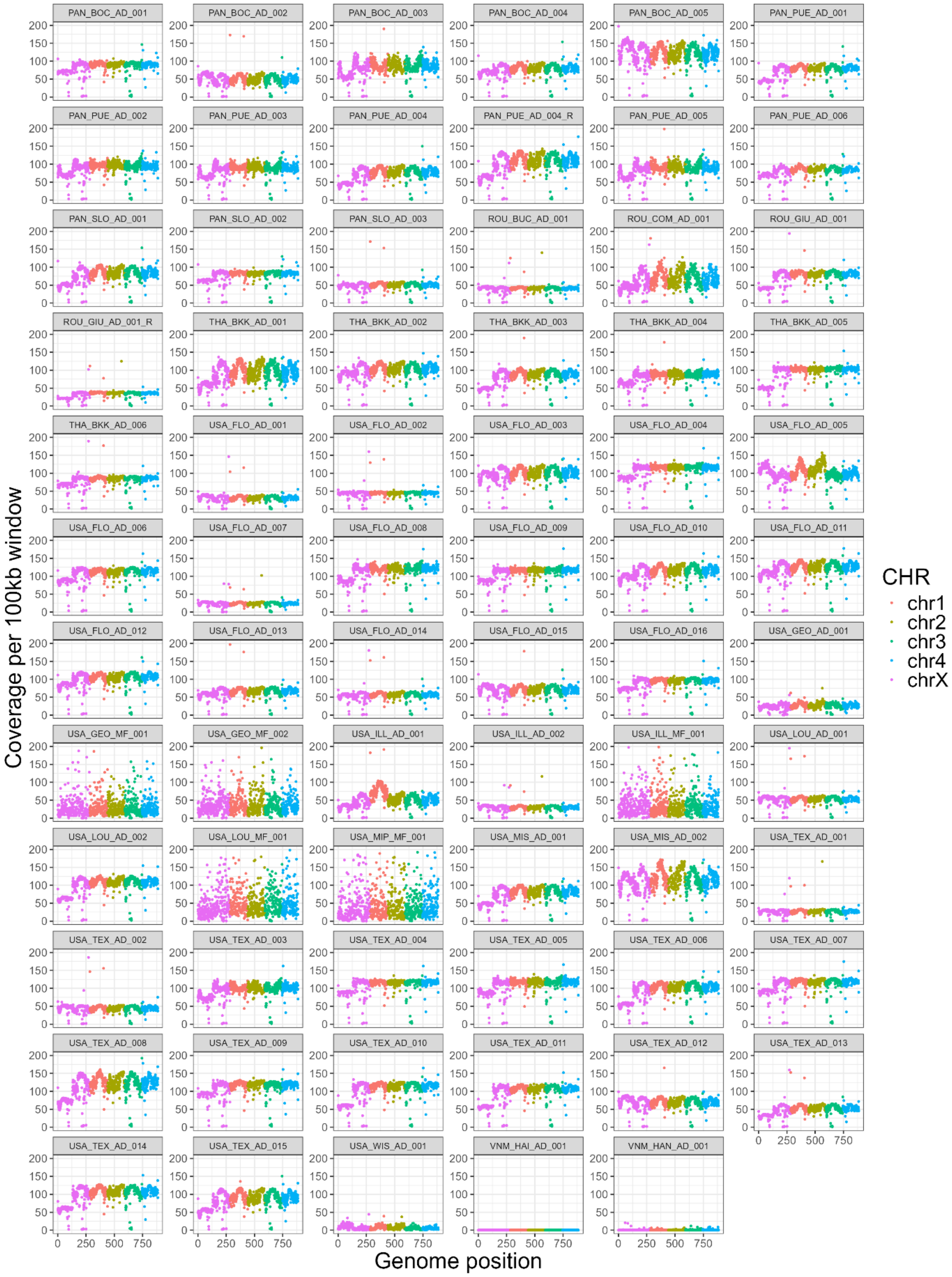


**Supplementary Fig. 2: Raw nuclear sequencing coverage per sample (Part 2/2).**

Coverage was calculated using 100 kb sliding windows for all mapped samples. Each chromosome is shown in a different colour, with a legend on the right.

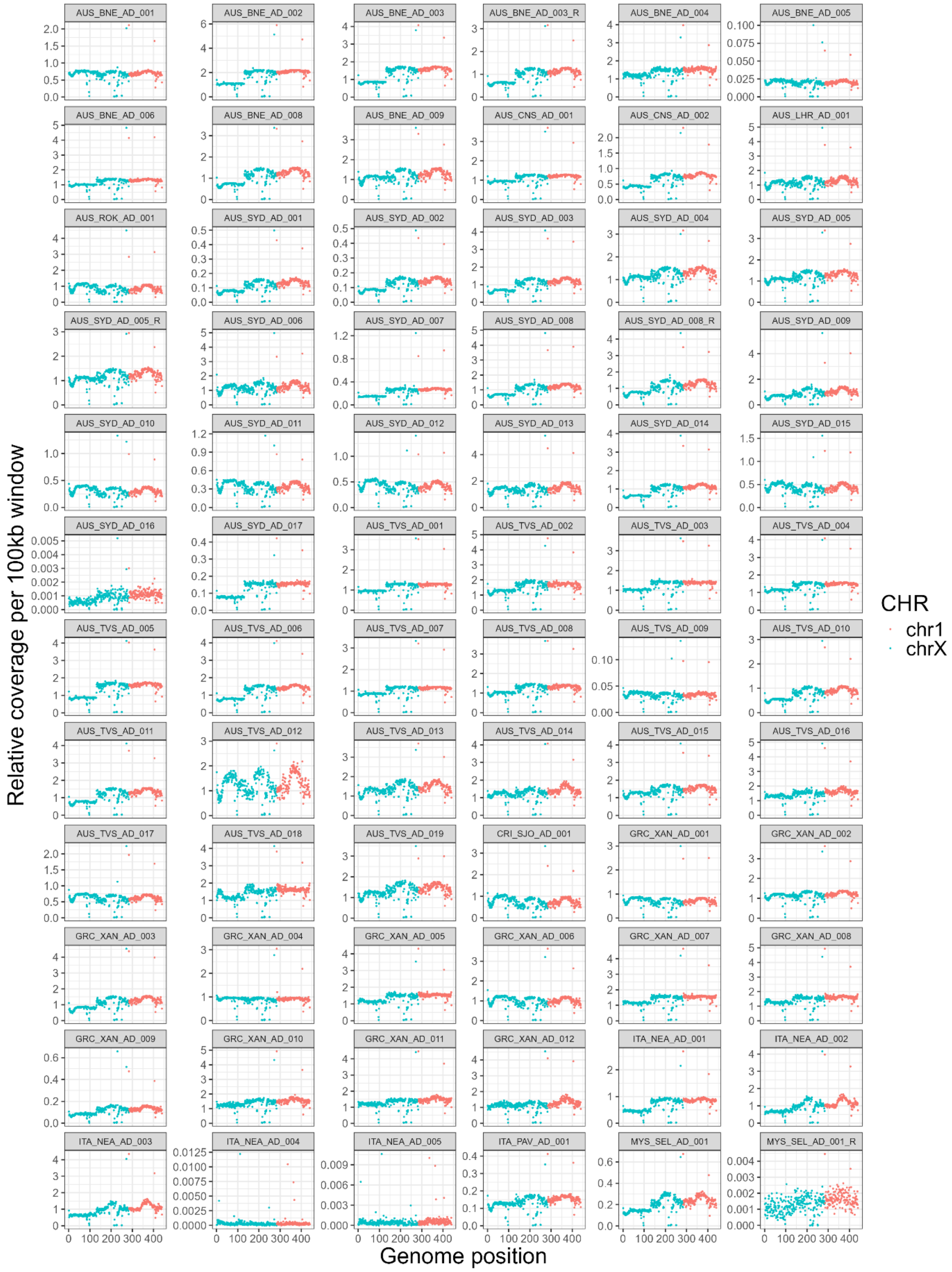


**Supplementary Fig. 3: Sex determination of samples based on relative coverage of sex-linked vs autosomal chromosomes (Part 1/2).**

Sequencing coverage was assessed using a 100kb sliding window and compared to the overall median coverage. Coverage of the sex-linked X chromosome (blue) was compared to an autosome (chromosome 1, red) to determine the parasite's sex. Male canine heartworms show reduced coverage on chromosome X (ratio ~0.5), while females show similar coverage across sex-linked and autosomal chromosomes (ratio ~1). Ambiguous results were observed in samples with low coverage. The remaining samples are shown in Supplementary Fig. 4.

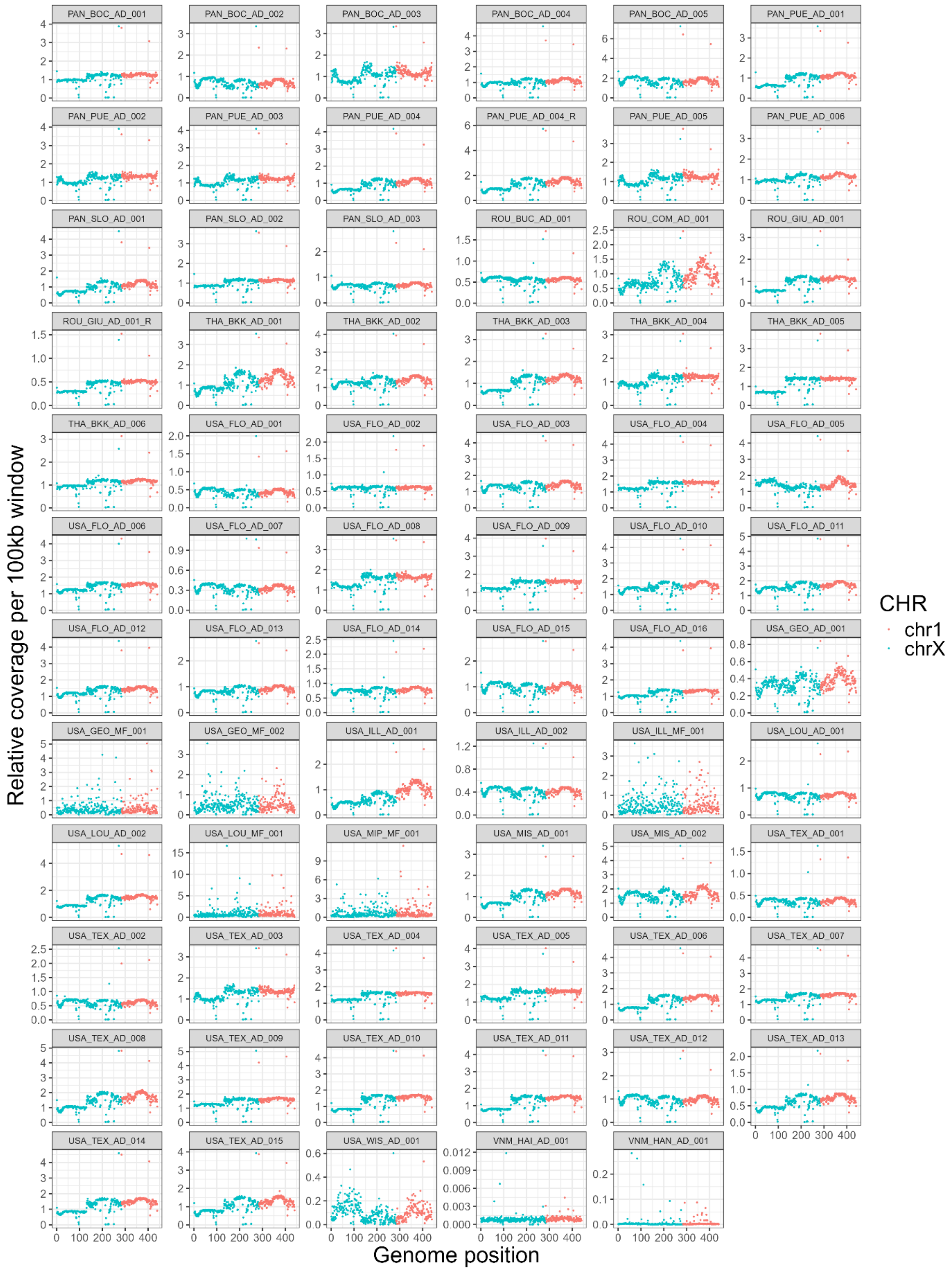


**Supplementary Fig. 4: Sex determination of samples based on relative coverage of sex-linked vs autosomal chromosomes (Part 2/2).**

Sequencing coverage was assessed using a 100kb sliding window and compared to the overall median coverage. Coverage of the sex-linked X chromosome (blue) was compared to an autosome (chromosome 1, red) to determine the parasite’s sex. Male canine heartworms show reduced coverage on chromosome X (ratio ~0.5), while females show similar coverage across sex-linked and autosomal chromosomes (ratio ~1). Ambiguous results were observed in samples with low coverage.

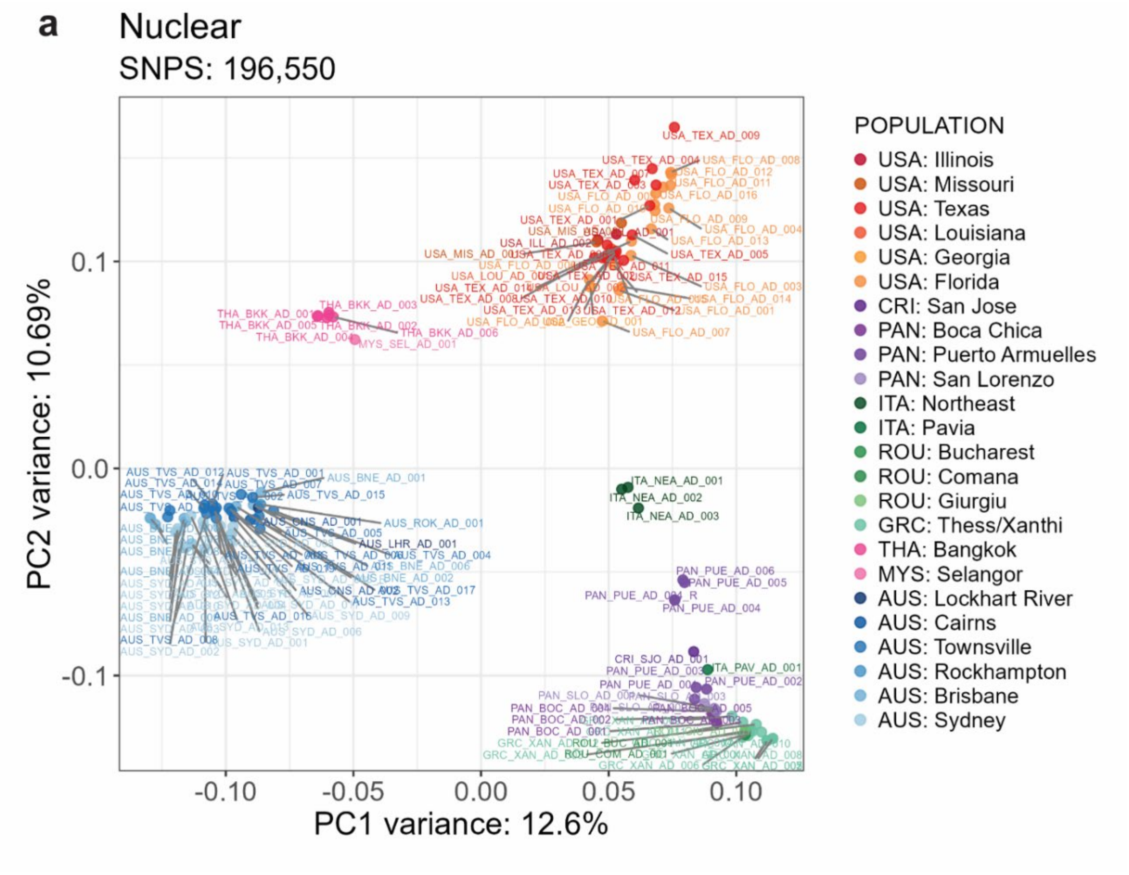


**Supplementary Fig. 5: Principal component analysis (PCA) of 196,550 nuclear single nucleotide polymorphisms (SNPs) from 124 heartworms with labelled metadata.**

Samples were collected from Australia (AUS), the USA, Panama (PAN), Greece (GRC), Thailand (THA), Italy (ITA), Romania (ROU), Costa Rica (CRI), and Malaysia (MYS). Replicate samples were included in the analyses. The legend on the right indicates the geographic origin of samples (i.e. COUNTRY: city or region). Principal component 1 (PC1) and principal component 2 (PC2) are shown on the x and y axes, explaining 12.6% and 10.69% of the variation, respectively. These figures are identical to Fig. 1b but include sample labels.

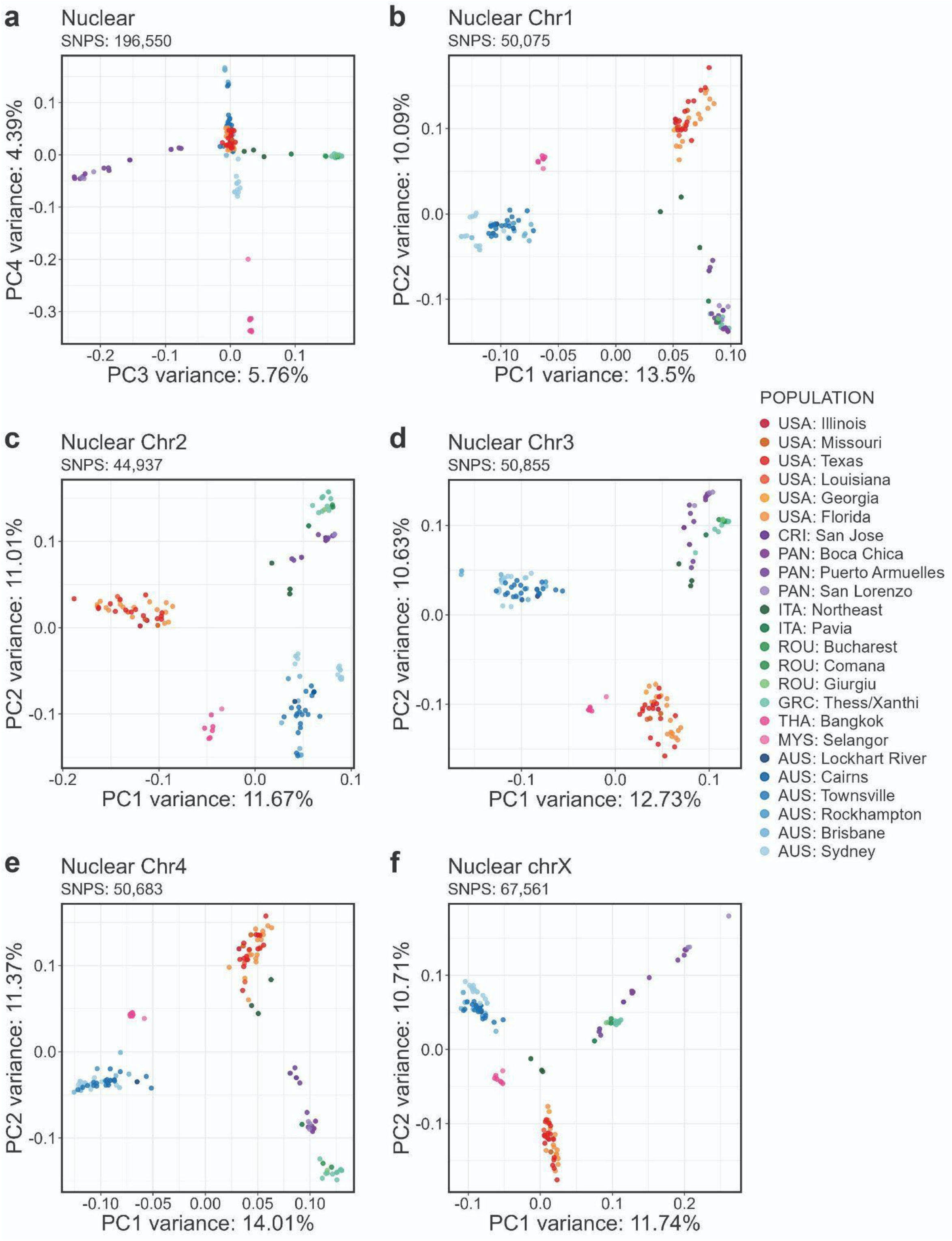


**Supplementary Fig. 6: Principal component analysis (PCA) of 124 heartworms from around the world based on the nuclear genome.**

Samples were collected from Australia (AUS), the USA, Panama (PAN), Greece (GRC), Thailand (THA), Italy (ITA), Romania (ROU), Costa Rica (CRI), and Malaysia (MYS). Replicate samples were included in the analyses. The legend on the right indicates the geographic origin of samples (i.e. COUNTRY: city or region). The principal components (PCs) are shown on the x and y axes. **a,** PCA of 196,550 nuclear single nucleotide polymorphisms (SNPs) showing PC3 and PC4, which explained 5.76% and 4.39% of the variation, respectively. **b–f,** PCAs of 44,937 – 67,561 SNPs in chromosome 1, chromosome 2, chromosome 3, chromosome 4, and the sex-linked X-chromosome of canine heartworm. PC1 and PC2 explained 11.67 – 14.01% and 10.09 – 11.37% of the variation, respectively.

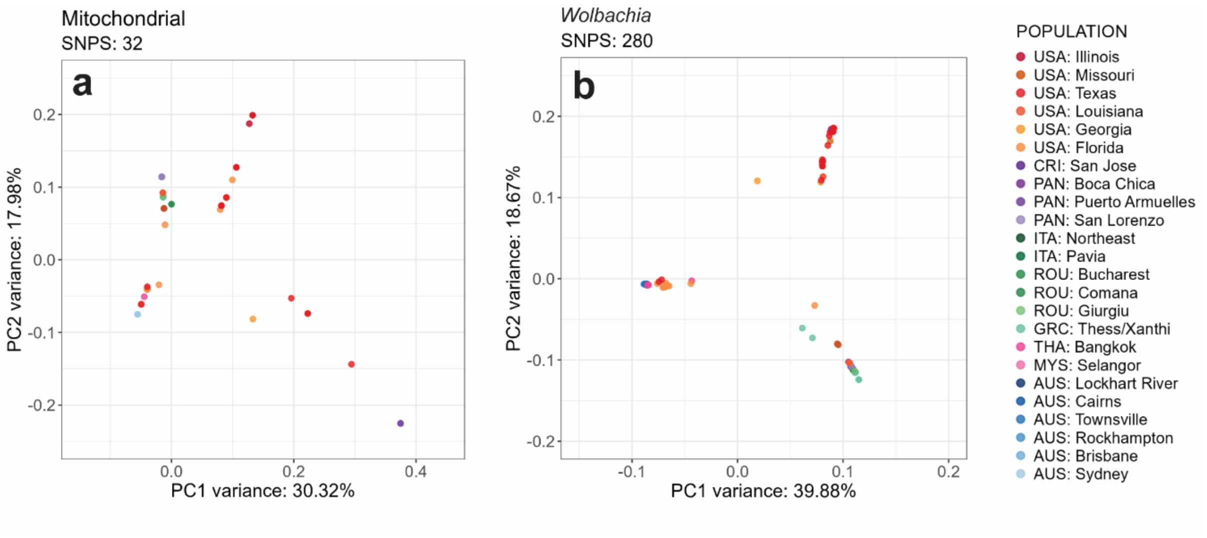


**Supplementary Fig. 7: Genetic relatedness of the mitochondria and *Wolbachia* endosymbionts of heartworms.**

**a,** Principal component analysis (PCA) of 32 mitochondrial single-nucleotide polymorphisms (SNPs) from 127 heartworms. Principal component 1 (PC1) and principal component 2 (PC2) explained 30.32% and 17.98% of the variation, respectively. **b,** PCA of 280 *Wolbachia* endosymbiont SNPs from 121 samples. PC1 and PC2 explained 39.88% and 18.67% of the variation, respectively. Replicate samples were included in all PCAs. The legend on the right indicates the geographic origin of samples (i.e. COUNTRY: city or region). Abbreviations: USA = United States of America; CRI = Costa Rica; PAN = Panama; ITA = Italy; ROU = Romania; GRC = Greece; VNM = Vietnam; THA = Thailand; MYS = Malaysia; AUS = Australia.

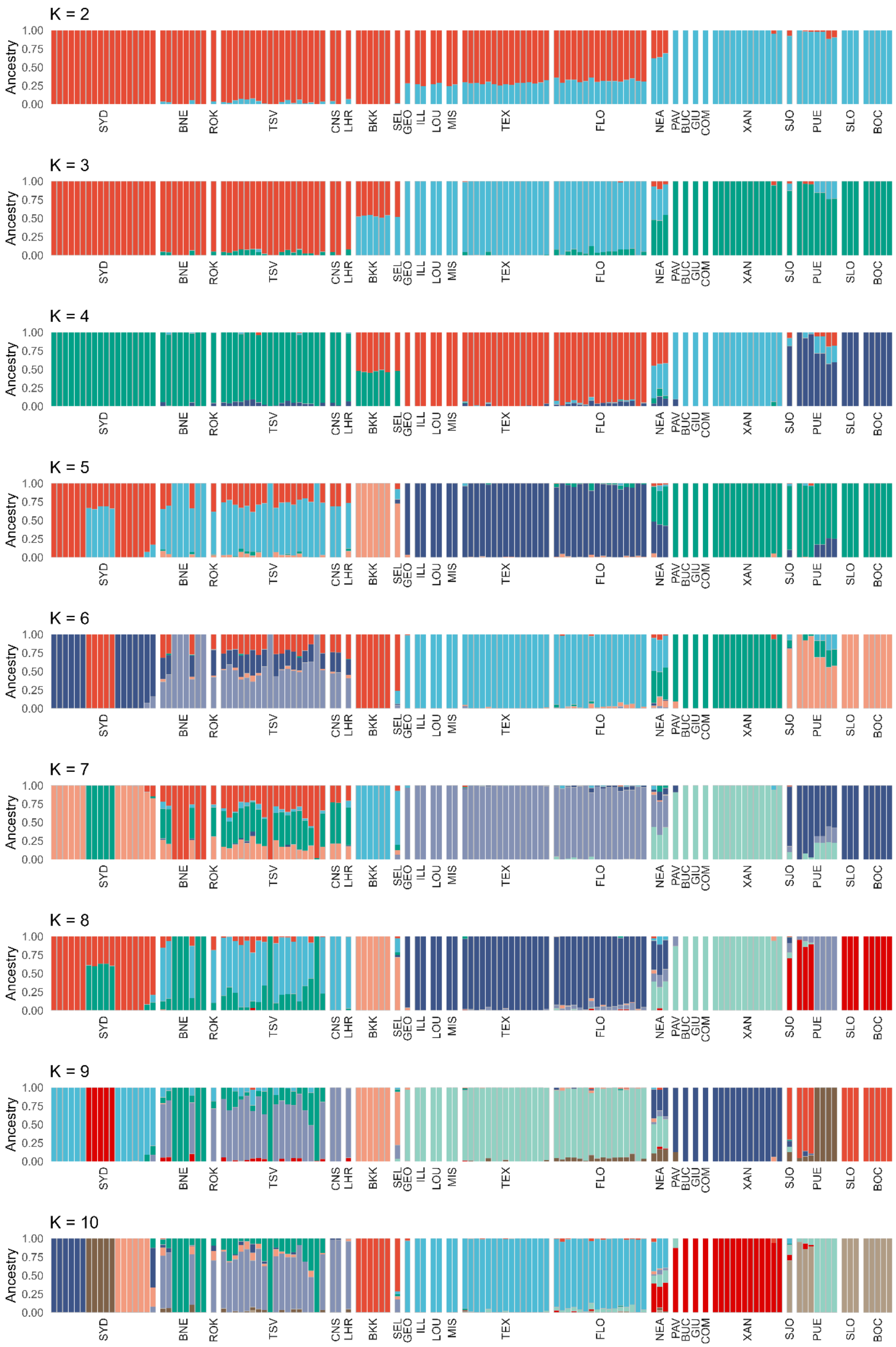

K = 2
K = 3
K = 4
K = 5
K = 6
K = 7
K = 8
K = 9
K = 10
Ancestry
1.00
0.75
0.50
0.25
0.00
SYD
BNE
ROK
TSV
CNS
LHR
BKK
SEL
GEO
ILL
LOU
MIS
TEX
FLO
NEA
PAV
BUC
GIU
COM
XAN
SJO
PUE
SLO
BOC

**Supplementary Fig. 8: Admixture analysis of adult heartworms.**

Analysis was performed in NGSadmix across a range of K values (2 – 10) (colours) for 124 heartworms (vertical bars). Geographic origins are indicated at the bottom of each chart. Abbreviations: Australia (Sydney: SYD, Brisbane: BNE, Rockhampton: ROK, Townsville: TSV, Cairns: CNS, Lockhart River: LHR); Thailand (Bangkok: BKK); Malaysia (Selangor: SEL); USA (Georgia: GEO, Illinois: ILL, Louisiana: LOU, Missouri: MIS, Texas: TEX, Florida: FLO); Italy (Northeast: NEA, Pavia: PAV); Romania (Bucharest: BUC, Giurgiu: GIU, Comana: COM); Greece (Thessaloniki/Xanthi: XAN); Costa Rica (San José: SJO); Panama (Puerto Armuelles: PUE, San Lorenzo: SLO, Boca Chica: BOC).

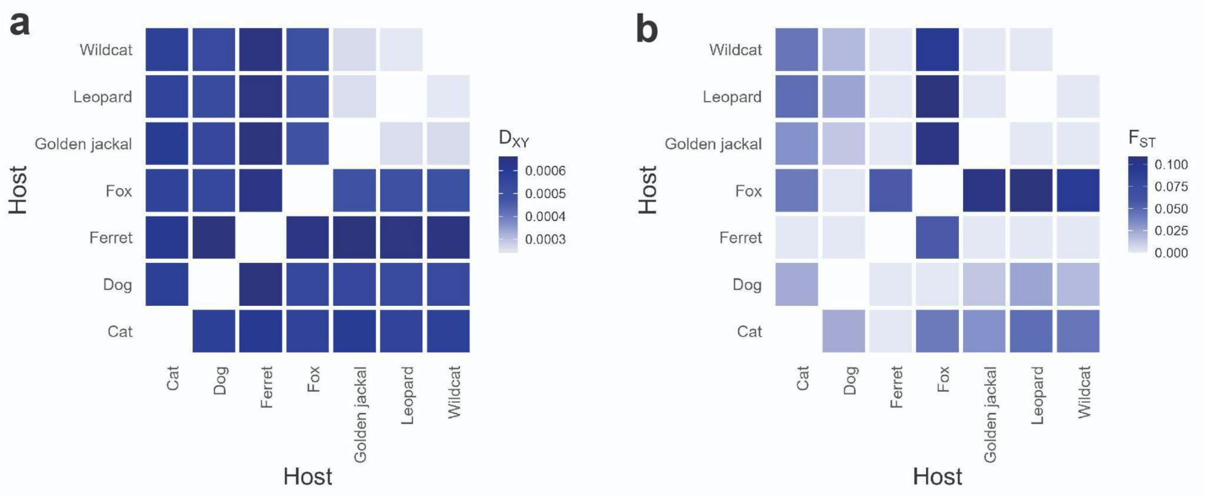


**Supplementary Fig. 9: Population genetic summary statistics of heartworms collected from diverse host populations.**

Heatmaps show **a,** Absolute nucleotide divergence ($D_{XY}$) and **b,** Genetic differentiation ($F_{ST}$) using the median of 100 kb sliding windows.

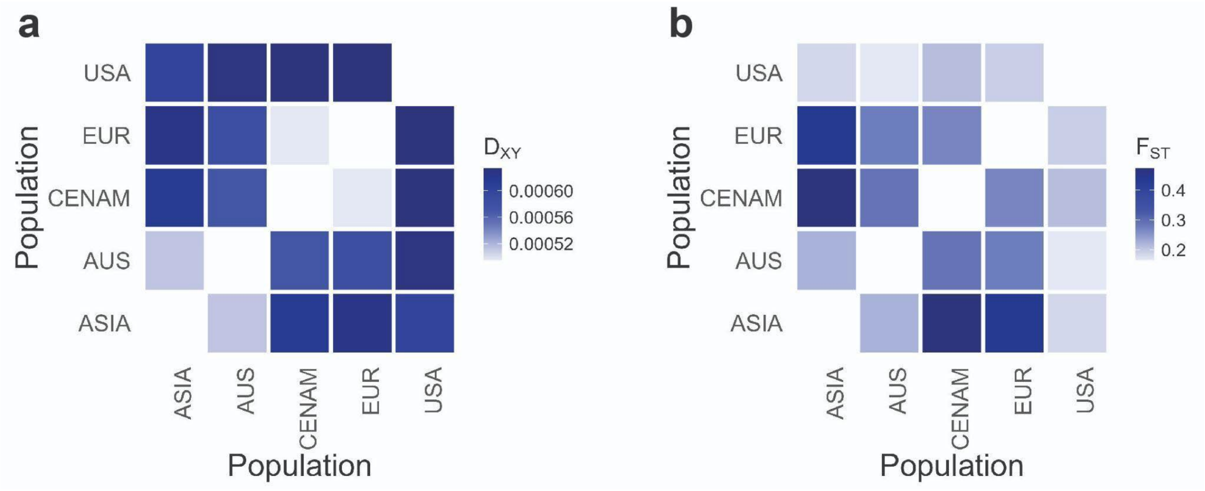


**Supplementary Fig. 10: Population genetic summary statistics of heartworms between broad geographical regions.**

Heatmaps show **a,** Absolute nucleotide divergence ($D_{XY}$) and **b,** Genetic differentiation ($F_{ST}$) using a 100 kb sliding window. Region abbreviations: ASIA = Asia; AUS = Australia; CENAM = Central America; EUR = Europe; USA = United States of America.

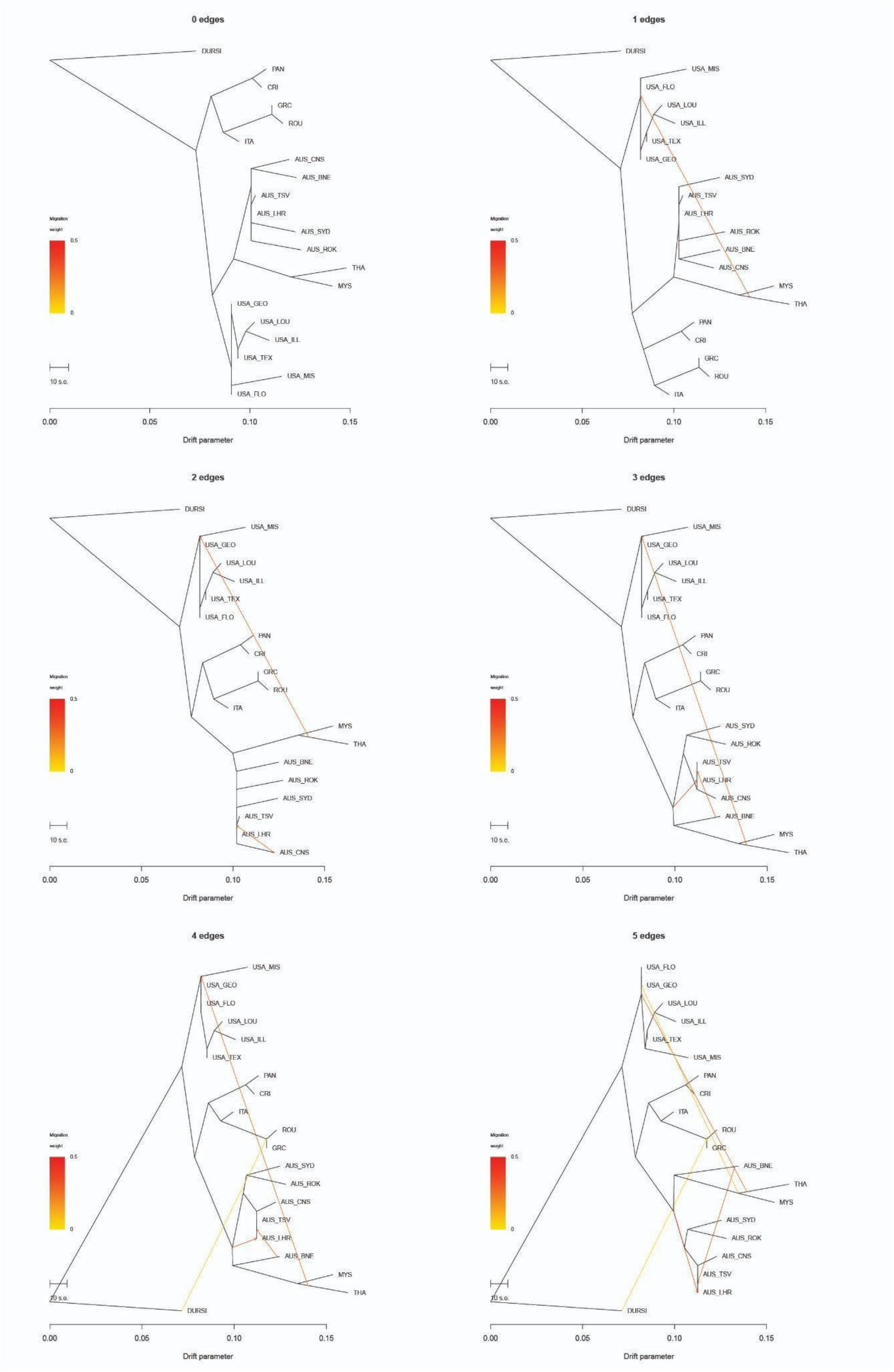
0 edges
DURSI
PAN
CRI
GRC
ROU
ITA
AUS_CNS
AUS_BNE
AUS_TSV
AUS_LHR
AUS_SYD
AUS_ROK
THA
MYS
USA_GEO
USA_LOU
USA_ILL
USA_TEX
USA_MIS
USA_FLO
Migration weight
0.5
0
10 s.e.
0.00
0.05
0.10
0.15
Drift parameter
1 edges
DURSI
USA_MIS
USA_FLO
USA_LOU
USA_ILL
USA_TEX
USA_GEO
AUS_SYD
AUS_TSV
AUS_LHR
AUS_ROK
AUS_BNE
AUS_CNS
MYS
THA
PAN
CRI
GRC
ROU
ITA
Migration weight
0.5
0
10 s.e.
0.00
0.05
0.10
0.15
Drift parameter
2 edges
DURSI
USA_MIS
USA_GEO
USA_LOU
USA_ILL
USA_TEX
USA_FLO
PAN
CRI
GRC
ROU
ITA
MYS
THA
AUS_BNE
AUS_ROK
AUS_SYD
AUS_TSV
AUS_LHR
AUS_CNS
Migration weight
0.5
0
10 s.e.
0.00
0.05
0.10
0.15
Drift parameter
3 edges
DURSI
USA_MIS
USA_GEO
USA_LOU
USA_ILL
USA_TEX
USA_FLO
PAN
CRI
GRC
ROU
ITA
AUS_SYD
AUS_ROK
AUS_TSV
AUS_LHR
AUS_CNS
AUS_BNE
MYS
THA
Migration weight
0.5
0
10 s.e.
0.00
0.05
0.10
0.15
Drift parameter
4 edges
USA_MIS
USA_GEO
USA_FLO
USA_LOU
USA_ILL
USA_TEX
PAN
CRI
ITA
ROU
GRC
AUS_SYD
AUS_ROK
AUS_CNS
AUS_TSV
AUS_LHR
AUS_BNE
MYS
THA
DURSI
Migration weight
0.5
0
10 s.e.
0.00
0.05
0.10
0.15
Drift parameter
5 edges
USA_FLO
USA_GEO
USA_LOU
USA_ILL
USA_TEX
USA_MIS
PAN
CRI
ITA
ROU
GRC
AUS_BNE
THA
MYS
AUS_SYD
AUS_ROK
AUS_CNS
AUS_TSV
AUS_LHR
DURSI
Migration weight
0.5
0
10 s.e.
0.00
0.05
0.10
0.15
Drift parameter

**Supplementary Fig. 11: TreeMix analysis of nuclear heartworm data.**

Maximum likelihood trees are shown with migration edges ranging from 0 to 5, using *Dirofilaria ursi* as an outgroup. Coloured lines represent inferred migration events, with the colour indicating support levels. Abbreviations: Australia: AUS (Sydney: SYD, Brisbane: BNE, Rockhampton: ROK, Townsville: TSV, Cairns: CNS, Lockhart River: LHR); USA (Georgia: GEO, Illinois: ILL, Louisiana: LOU, Missouri: MIS, Texas: TEX, Florida: FLO); Thailand: THA; Malaysia: MYS; Italy: ITA; Romania: ROU; Greece: GRC; Costa Rica: CRI; Panama: PAN.

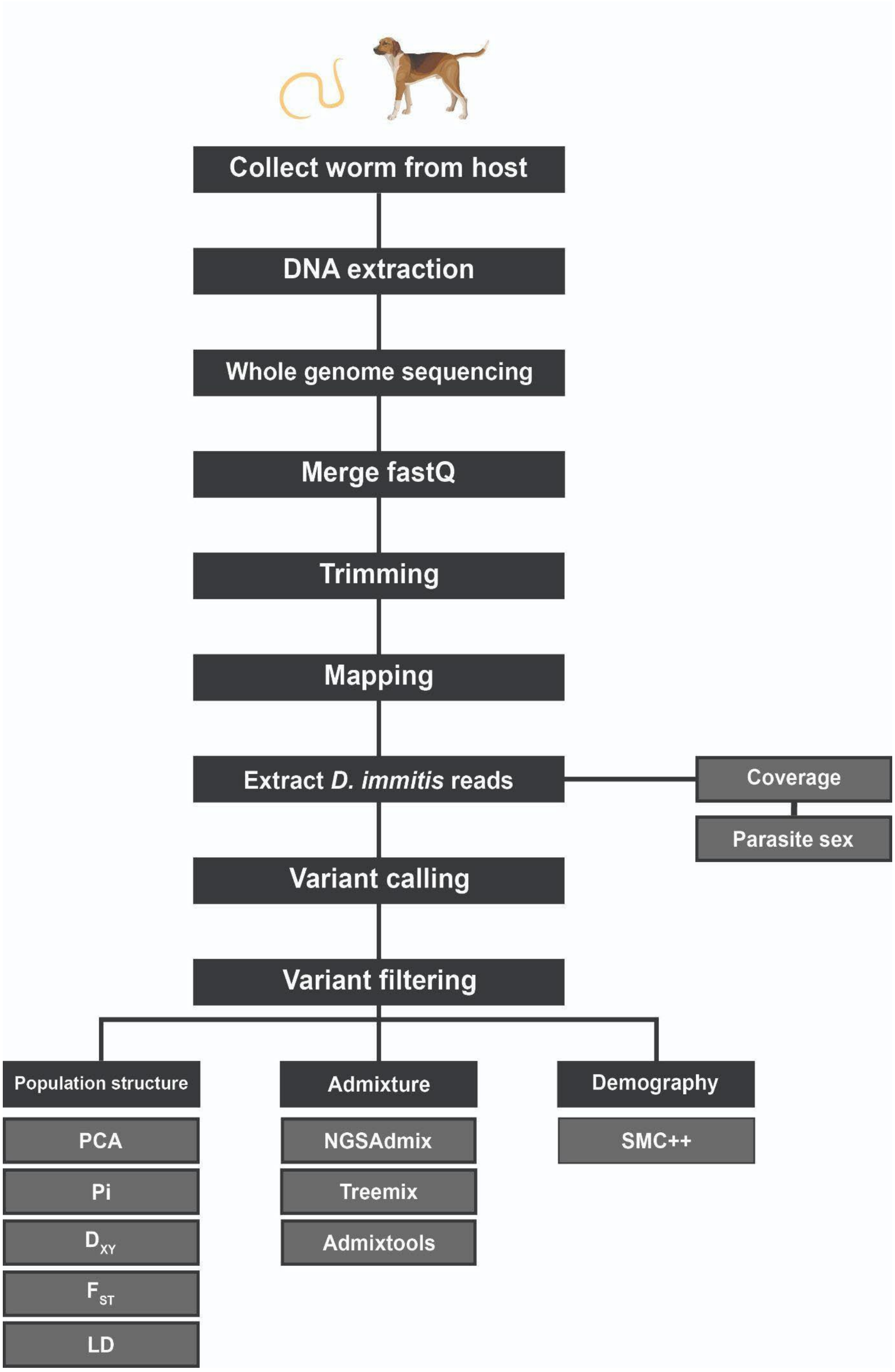


**Supplementary Fig. 12: Summary diagram of the population genomics workflow in this study.**

Abbreviations of population structure analyses: PCA = Principal Component Analysis; Pi = nucleotide diversity; $D_{XY}$ = absolute nucleotide divergence; $F_{ST}$ = genetic differentiation.

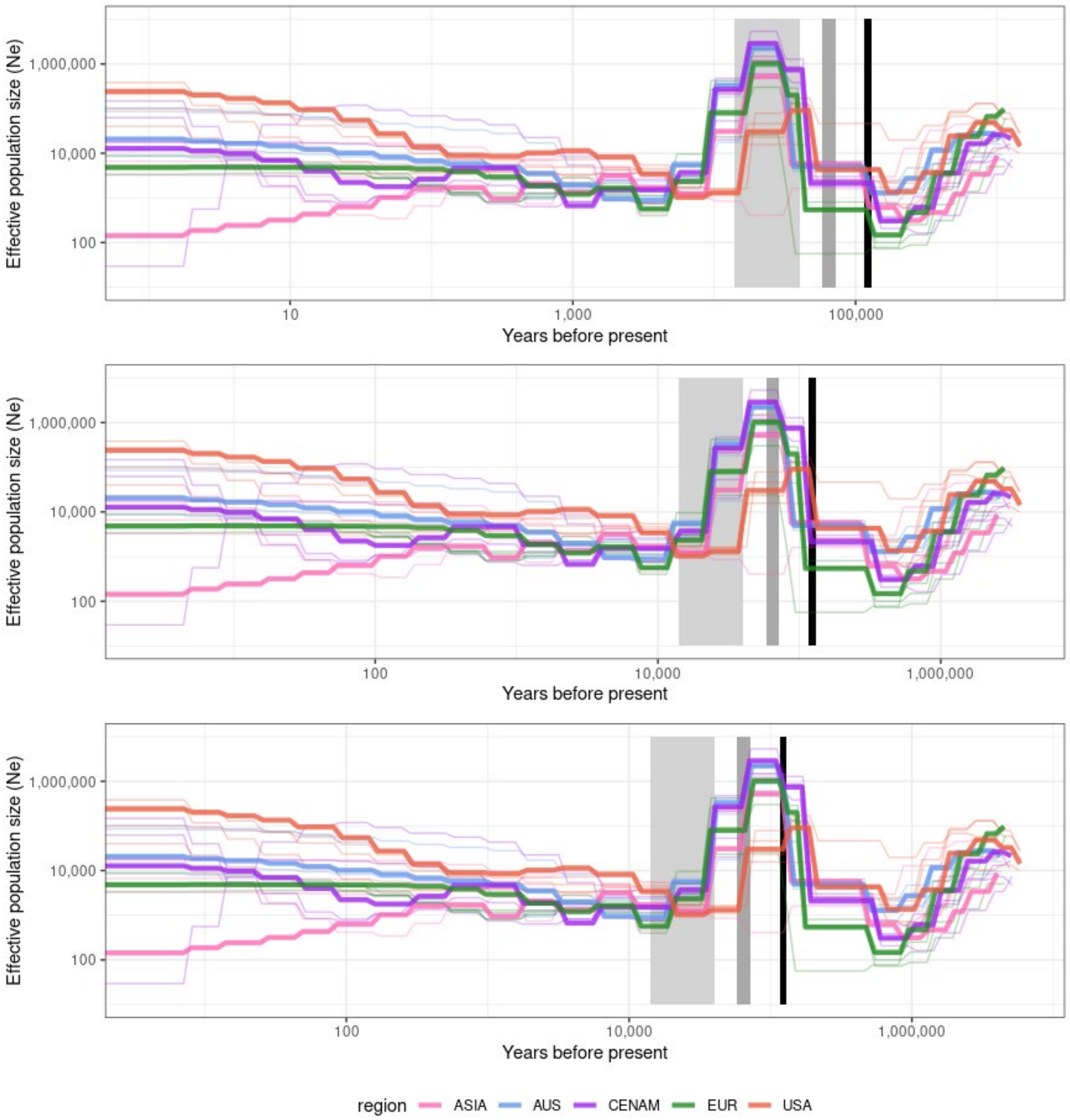


**Supplementary Fig. 13: Demographic history analysis of heartworms from dogs using a range of generation times.**

Size histories were inferred using SMC++, with each line representing a heartworm subpopulation. A range of generation times were tested for *Dirofilaria immitis* (i.e. 1, 2.5, and 4 years). The grey box indicates previously suggested date ranges for dog domestication (~14–40 kya). Abbreviations: AUS = Australia; CENAM = Central America; EUR = Europe; USA = United States of America; MIS 4 = Marine Isotype Stage 4.

### Supplementary Table 1: Adult heartworm burdens in Carnivora hosts

| Host | Location | Animals | Adult worms /animal | Reference |
|---|---|---|---|---|
| **Wolves** | | | | |
| Gray wolves | Italy | 3 | 5 | Moroni et al., 2020 |
| Wolf (*Canis lupus*) | Serbia | 1 | 37 | Penezić et al., 2014 |
| Wolf (*Canis lupus*) | Spain | 1 | 1 | Segovia et al., 2001 |
| Grey wolf (*Canis lupus lupus*) | Serbia | 1 | 42 | Gavrilović et al., 2015 |
| Red wolves (*Canis rufus gregoryi*) | USA | 8 | 77.8 | Custer & Pence, 1981 |
| **Microfilaremia reported?** | Yes | | | Gomes-de-Sá et al., 2022 |
| **Coyotes** | | | | |
| Coyotes (*Canis latrans*) | USA | 77 | 21.29 | Aher et al., 2016 |
| Coyotes (*Canis latrans*) | USA | 36 | 12 | King & Bohning, 1984 |
| Coyotes (*Canis latrans*) | USA | 2 | 3 | Wixsom et al., 1991 |
| Coyotes (*Canis latrans*) | USA | 17 | 13.2 | Wixsom et al., 1991 |
| Coyote (Canis latrans) | USA | 1 | 11 | Kazacos & Edberg, 1979 |
| Coyotes (*Canis latrans*) | USA | 21 | 19.4 | Sacks, 1998 |
| Coyotes (*Canis latrans*) | USA | 147 | 8.7 | Nelson et al., 2003 |
| Coyotes (*Canis latrans*) | USA | 8 | 13.6 | Agostine & Jones, 1982 |
| Coyotes (*Canis latrans*) | USA | 1 | 5 | Agostine & Jones, 1982 |
| Coyotes (*Canis latrans*) | USA | 1 | 3 | Agostine & Jones, 1982 |
| Coyotes (*Canis latrans*) | USA | 20 | 9 | Weinmann & Garcia, 1980 |
| Coyotes (*Canis latrans*) | USA | 19 | 16.2 | Weinmann & Garcia, 1980 |
| Coyotes (*Canis latrans*) | USA | 17 | 13.6 | Custer & Pence, 1981 |
| **Microfilaremia reported?** | Yes | | | Weinmann & Garcia, 1980 |
| **Jackals** | | | | |
| Eurasian/golden jackals (*Canis aureus*) | Iran | 4 | 5.25 | Heidari et al., 2015 |
| Golden jackals (*Canis aureus*) | Iran | 8 | 6.62 | Sharifdini et al., 2022 |

| | | | | |
|---|---|---|---|---|
| Golden jackals (*Canis aureus*) | Bulgaria | 122 | 4.1 | Panayotova-Pencheva et al., 2016 |
| Jackals (*Canis aureus*) | Russia | 14 | 12 | Kravchenko et al., 2016 |
| Golden jackals (*Canis aureus*) | Romania | 12 | 2.92 | Ionică et al., 2022 |
| Golden jackals (*Canis aureus*) | Romania | 10 | 3 | Ionică et al., 2016 |
| Golden jackals (*Canis aureus*) | Hungary | 2 | 1 | Tolnai et al., 2014 |
| **Microfilaremia reported?** | Yes | | | Ionică et al., 2016 |
| **Foxes** | | | | |
| Red foxes (*Vulpes vulpes*) | Spain | 46 | 4.39 | Gortazar et al., 1994 |
| Red foxes (*Vulpes vulpes*) | Spain | N/A | 4.4 | Gortázar et al., 1998 |
| Gray fox (*Urocyon cinereoargenteus*) | USA | 2 | 7 | Simmons et al., 1980 |
| Gray fox (*Urocyon cinereoargenteus*) | USA | 3 | 2 | King & Bohning, 1984 |
| Red fox (*Vulpes vulpes*) | USA | 1 | 5 | King & Bohning, 1984 |
| Red foxes (*Vulpes vulpes*) | USA | 5 | 3 | Wixsom et al., 1991 |
| Red foxes (*Vulpes fulva*) & Gray foxes (*Urocyon cinereoargenteus*) | USA | 6 | 5 | Kazacos & Edberg, 1979 |
| Red foxes (*Vulpes vulpes L.*) | Bulgaria | 29 | 4.79 | Panayotova-Pencheva et al., 2016 |
| Foxes (*Vulpes vulpes*) | Russia | 48 | 9.2 | Kravchenko et al., 2016 |
| Red foxes (*Vulpes vulpes*) | Romania | 4 | 1.5 | Ionică et al., 2022 |
| Red Fox (*Vulpes vulpes*) | Hungary | 20 | 1.5 | Tolnai et al., 2014 |
| **Microfilaremia reported?** | Yes, but there is low risk that they are a competent reservoir. | | | Marks & Bloomfield, 1998<br>McCall et al., 2008 |
| **Felidae** | | | | |
| Wild cat (*Felis silvestris*) | Serbia | 1 | 2 | Penezić et al., 2014 |
| Wild cats (*Felis silvestris*) | Romania | 2 | 1 | Ionică et al., 2022 |
| Cats | USA | 2 | 1.5 | Nelson & Johnson, 2024 |

| | | | | |
|---|---|---|---|---|
| Snow leopard (*Uncia uncia*) | Japan | 1 | 3 | Murata et al., 2003 |
| **Microfilaremia reported?** | Infrequently, not considered competent reservoir. | | | McCall et al., 2008 |
| **Procyonidae** | | | | |
| Raccoon dogs (*Nyctereutes procyonoides*) | Russia | 28 | 12.6 | Kravchenko et al., 2016 |
| Raccoon dog (*Nyctereutes procyonoides*) | Romania | 1 | 1 | Ionică et al., 2022 |
| Raccoon dogs (*Nyctereutes procyonoides viverrinus*) | Japan | 8 | 1.75 | Nakagaki et al., 2000 |
| **Microfilaremia reported?** | Infrequently, not considered competent reservoir. | | | Ionică et al., 2022 |
| **Mustelidae** | | | | |
| European badger (*Meles meles*) | Romania | 1 | 3 | Ionică et al., 2022 |
| Ferret (*Mustela putorius furo*) | Hungary | 1 | 2 | Molnár et al., 2010 |
| European badgers (*Meles meles*) | Greece | 2 | 3 | Markakis et al., 2024 |
| **Microfilaremia reported?** | Infrequently, not considered competent reservoir. | | | McCall et al., 2008 |
| **Pinnipedia** | | | | |
| Seal (*Phoca vitulina*) | Portugal (zoo) | 1 | 32 | Alho et al., 2017 |
| Seal (*Arctocephalus pusillus pusillus*) | Portugal (zoo) | 3 | 17 | Alho et al., 2017 |
| Harbor seal (*Phoca vitulina*) | South Korea (zoo) | 1 | 2 | Kang et al., 2002 |
| **Microfilaremia reported?** | Infrequently, not considered competent reservoir. | | | Alho et al., 2017 |
| **Ursidae** | | | | |
| Brown bear (*Ursus arctos*) | Greece | 1 | 4 | Papadopoulos et al., 2017 |
| Black bear (*Ursus americanus*) | USA | 4 | 3 | Crum et al., 1978 |
| Black bear (*Ursus americanus*) | USA | 1 | 5 | Johnson, 1975 |
| **Microfilaremia reported?** | Infrequently, not considered competent reservoir. | | | McCall et al., 2008 |
| **Dogs** | | | | |
| Dogs | Iran | 10 | 23.5 | Sharifdini et al., 2022 |
| Dogs | Bulgaria | 9 | 14.43 | Panayotova-Pencheva et al., 2016 |
| Dog | USA | 1 | 150 | Oliveira et al., 2021 |
| Dogs | Italy | 2 | 15.5 | Santoro et al., 2019 |

| | | | | |
|---|---|---|---|---|
| Dogs | Taiwan | 477 | 7.2 | Wu & Fan, 2003 |
| Dogs | Mexico | 52 | 4.5 | Bolio-Gonzalez et al., 2007 |
| Dogs | USA | 170 | 14 | Kaiser & Williams, 2004 |
| Dogs | Australia | 15 | 5.8 | Bidgood & Collins, 1996 |
| Dogs | Hungary | 2 | 9 | Tolnai et al., 2014 |
| Dogs | N/A | 6 | 23.83 | Rafailov et al., 2022 |
| Dogs | USA | 50 | 7.62 | Henry et al., 2018 |
| Dogs | USA | 50 | 24.62 | Henry et al., 2018 |
| Dogs | USA | 50 | 53.86 | Henry et al., 2018 |
| Dogs | USA | 50 | 104.92 | Henry et al., 2018 |
| Dogs | Panama | 3 | 44 | Chacón & Candanedo, 2021 |
| **Microfilaremia reported?** | Yes | | | Panetta et al., 2021 |
