## Supplementary Data 1 for "Population genomics reveals an ancient origin of heartworms in canids"

**Supplementary Data 1: Genome mapping statistics and coverage for nuclear, mitochondrial, and *Wolbachia* genomes.**

Abbreviations: DI = *Dirofilaria immitis* ; REP = Replicate sample; M = Millions

| Sample ID | Original ID | Collection | Sample type | BAM Total Reads (M) | Mapped Reads (M) | % Mapped | Extracted DI Reads (M) | % Mapped to DI | Nuclear coverage (mean) | Nuclear coverage (sd) | Mitochondrial coverage (mean) | *Wolbachia* coverage (mean) |
|---|---|---|---|---|---|---|---|---|---|---|---|---|
| AUS_BNE_AD_001 | JS6342 | This study | DI | 71.7 | 71.3 | 99.4 | 71 | 99.0 | 48.39 | 10.86 | 13695.00 | 2915.19 |
| AUS_BNE_AD_002 | JS6343 | This study | DI | 128.6 | 127.9 | 99.5 | 127.5 | 99.1 | 133.68 | 37.73 | 20973.50 | 340.66 |
| AUS_BNE_AD_003 | JS6344 | This study | DI | 101.5 | 101.1 | 99.6 | 100.6 | 99.1 | 102.01 | 28.17 | 23016.90 | 777.32 |
| AUS_BNE_AD_003_R | JS6607 | This study | DI (REP) | 72.2 | 72 | 99.7 | 71.9 | 99.6 | 72.98 | 21.08 | 21823.20 | 1050.74 |
| AUS_BNE_AD_004 | JS6345 | This study | DI | 90.8 | 90.2 | 99.3 | 89.9 | 99.0 | 100.49 | 21.01 | 558.64 | 9.20 |
| AUS_BNE_AD_005 | JS6346 | This study | DI | 7.5 | 7.3 | 97.3 | 6.7 | 89.3 | 1.41 | 0.42 | 3936.33 | 575.08 |
| AUS_BNE_AD_006 | JS6347 | This study | DI | 80.9 | 80.4 | 99.4 | 79.3 | 98.0 | 87.60 | 24.55 | 3066.31 | 80.32 |
| AUS_BNE_AD_008 | JS6610 | This study | DI | 80.1 | 79.9 | 99.8 | 75.8 | 94.6 | 82.61 | 23.58 | 22138.70 | 349.02 |
| AUS_BNE_AD_009 | JS6611 | This study | DI | 74.2 | 74.1 | 99.9 | 74 | 99.7 | 87.03 | 21.85 | 2123.36 | 92.01 |
| AUS_CNS_AD_001 | JS6349 | This study | DI | 84.6 | 84.2 | 99.5 | 80.4 | 95.0 | 81.39 | 19.36 | 4884.56 | 946.81 |
| AUS_CNS_AD_002 | JS6350 | This study | DI | 72.5 | 72.2 | 99.6 | 70.8 | 97.7 | 49.53 | 14.26 | 10168.00 | 2828.42 |
| AUS_LHR_AD_001 | JS6281 | This study | DI | 74.6 | 74.3 | 99.6 | 74.1 | 99.3 | 87.67 | 26.54 | 2961.65 | 91.00 |
| AUS_ROK_AD_001 | JS6606 | This study | DI | 77.3 | 77.1 | 99.7 | 73.7 | 95.3 | 59.86 | 22.06 | 30692.50 | 1950.34 |
| AUS_SYD_AD_001 | JS5877 | Public data | DI | 8.5 | 8.5 | 100.0 | 8.4 | 98.8 | 8.78 | 2.89 | 3421.03 | 125.25 |
| AUS_SYD_AD_002 | JS5876 | Public data | DI | 8.4 | 8.4 | 100.0 | 8.2 | 97.6 | 9.45 | 2.97 | 4734.91 | 2.17 |
| AUS_SYD_AD_003 | JS5875 | Public data | DI | 65.3 | 65.2 | 99.8 | 65 | 99.5 | 78.05 | 24.76 | 12507.90 | 231.42 |
| AUS_SYD_AD_004 | JS5874 | Public data | DI | 71.5 | 71.3 | 99.7 | 71.1 | 99.4 | 89.25 | 20.31 | 1926.36 | 165.62 |
| AUS_SYD_AD_005 | JS5873 | Public data | DI | 69.8 | 69.7 | 99.9 | 69.5 | 99.6 | 87.88 | 20.02 | 6849.22 | 464.06 |
| AUS_SYD_AD_005_R | JS6609 | This study | DI (REP) | 72.8 | 72.7 | 99.9 | 72.6 | 99.7 | 84.84 | 19.21 | 5398.47 | 219.23 |
| AUS_SYD_AD_006 | JS6277 | This study | DI | 77.5 | 77.2 | 99.6 | 76.8 | 99.1 | 87.34 | 27.42 | 5108.39 | 218.72 |
| AUS_SYD_AD_007 | JS6278 | This study | DI | 76.5 | 47.8 | 62.5 | 47.4 | 62.0 | 17.44 | 6.35 | 1826.77 | 45.29 |
| AUS_SYD_AD_008 | JS6279 | This study | DI | 77 | 76.5 | 99.4 | 76.2 | 99.0 | 79.68 | 27.13 | 7534.26 | 751.86 |
| AUS_SYD_AD_008_R | JS6608 | This study | DI (REP) | 78.1 | 77.9 | 99.7 | 77.8 | 99.6 | 81.94 | 26.71 | 17456.00 | 695.59 |
| AUS_SYD_AD_009 | JS6280 | This study | DI | 70.6 | 69.7 | 98.7 | 69.5 | 98.4 | 69.48 | 29.45 | 4694.63 | 819.19 |
| AUS_SYD_AD_010 | JS6601 | This study | DI | 99.1 | 99 | 99.9 | 98.4 | 99.3 | 21.35 | 6.91 | 29404.90 | 7999.64 |
| AUS_SYD_AD_011 | JS6602 | This study | DI | 84.1 | 83.9 | 99.8 | 83.4 | 99.2 | 23.84 | 6.55 | 27070.50 | 6587.44 |
| AUS_SYD_AD_012 | JS6603 | This study | DI | 87.7 | 87.6 | 99.9 | 87.2 | 99.4 | 29.25 | 7.96 | 36777.60 | 6056.63 |
| AUS_SYD_AD_013 | JS6756 | This study | DI | 100.1 | 99.8 | 99.7 | 99.6 | 99.5 | 101.21 | 29.18 | 6003.53 | 1151.50 |
| AUS_SYD_AD_014 | JS6757 | This study | DI | 70.5 | 70.2 | 99.6 | 70 | 99.3 | 72.93 | 22.64 | 12142.60 | 818.83 |
| AUS_SYD_AD_015 | JS6758 | This study | DI | 78.2 | 77.9 | 99.6 | 77.2 | 98.7 | 30.20 | 8.53 | 21155.70 | 5191.85 |
| AUS_SYD_AD_016 | JS6765 | This study | DI | 0.5 | 0.5 | 100.0 | 0.5 | 100.0 | 0.07 | 0.03 | 226.74 | 35.97 |
| AUS_SYD_AD_017 | JS6766 | This study | DI | 29.4 | 29.1 | 99.0 | 25.4 | 86.4 | 9.94 | 2.81 | 4656.44 | 325.85 |
| AUS_TVS_AD_001 | JS6351 | This study | DI | 75.5 | 75.1 | 99.5 | 75 | 99.3 | 84.44 | 19.92 | 2297.54 | 63.43 |
| AUS_TVS_AD_002 | JS6352 | This study | DI | 109.7 | 109.1 | 99.5 | 108.8 | 99.2 | 114.68 | 27.53 | 4349.69 | 232.24 |
| AUS_TVS_AD_003 | JS6353 | This study | DI | 80.5 | 80.2 | 99.6 | 80 | 99.4 | 92.78 | 21.33 | 2463.58 | 50.82 |
| AUS_TVS_AD_004 | JS6354 | This study | DI | 88.7 | 88.3 | 99.5 | 88.1 | 99.3 | 99.47 | 23.06 | 2885.98 | 120.20 |
| AUS_TVS_AD_005 | JS6355 | This study | DI | 110.2 | 109.7 | 99.5 | 109.4 | 99.3 | 103.73 | 29.06 | 25195.60 | 961.92 |
| AUS_TVS_AD_006 | JS6356 | This study | DI | 93.3 | 92.8 | 99.5 | 92.5 | 99.1 | 92.76 | 26.67 | 19831.20 | 834.00 |
| AUS_TVS_AD_007 | JS6357 | This study | DI | 69.3 | 69 | 99.6 | 68.8 | 99.3 | 76.62 | 18.42 | 2212.00 | 59.89 |
| AUS_TVS_AD_008 | JS6358 | This study | DI | 81 | 80.5 | 99.4 | 80.3 | 99.1 | 88.99 | 21.43 | 2565.00 | 67.28 |
| AUS_TVS_AD_009 | JS6359 | This study | DI | 6.9 | 6.9 | 100.0 | 6.9 | 100.0 | 2.35 | 0.65 | 2766.60 | 520.26 |
| AUS_TVS_AD_010 | JS6360 | This study | DI | 66.4 | 65.7 | 98.9 | 65.5 | 98.6 | 59.43 | 18.13 | 21505.00 | 1373.55 |
| AUS_TVS_AD_011 | JS6368 | This study | DI | 84 | 83.8 | 99.8 | 83.5 | 99.4 | 87.20 | 26.26 | 23694.80 | 883.82 |
| AUS_TVS_AD_012 | JS6369 | This study | DI | 85.5 | 85.2 | 99.6 | 85 | 99.4 | 88.20 | 29.08 | 3022.42 | 146.37 |
| AUS_TVS_AD_013 | JS6370 | This study | DI | 88 | 87.8 | 99.8 | 87.6 | 99.5 | 101.09 | 24.43 | 5868.40 | 177.28 |
| AUS_TVS_AD_014 | JS6612 | This study | DI | 81.3 | 81.1 | 99.8 | 80.9 | 99.5 | 93.31 | 23.30 | 6379.35 | 141.65 |
| AUS_TVS_AD_015 | JS6613 | This study | DI | 89 | 88.8 | 99.8 | 88.6 | 99.6 | 102.48 | 23.95 | 3106.15 | 140.66 |
| AUS_TVS_AD_016 | JS6614 | This study | DI | 91.7 | 91.4 | 99.7 | 91.2 | 99.5 | 106.19 | 26.33 | 7349.46 | 18.16 |
| AUS_TVS_AD_017 | JS6615 | This study | DI | 85.3 | 85.2 | 99.9 | 85 | 99.6 | 43.79 | 11.38 | 29200.00 | 5031.36 |
| AUS_TVS_AD_018 | JS6616 | This study | DI | 94.8 | 94.5 | 99.7 | 94.3 | 99.5 | 106.77 | 24.92 | 4185.09 | 42.24 |
| AUS_TVS_AD_019 | JS6617 | This study | DI | 86.4 | 86.2 | 99.8 | 86.1 | 99.7 | 98.03 | 23.49 | 3675.94 | 158.40 |
| CRI_SJO_AD_001 | JS6670 | This study | DI | 66.2 | 65.8 | 99.4 | 53.3 | 80.5 | 52.20 | 17.60 | 20931.30 | 343.98 |
| GRC_XAN_AD_001 | JS6734 | This study | DI | 72.8 | 72.1 | 99.0 | 67.2 | 92.3 | 50.17 | 14.62 | 15891.30 | 1638.22 |
| GRC_XAN_AD_002 | JS6735 | This study | DI | 81.2 | 80.8 | 99.5 | 80.4 | 99.0 | 83.32 | 18.55 | 15025.80 | 1255.28 |
| GRC_XAN_AD_003 | JS6736 | This study | DI | 75.1 | 74.6 | 99.3 | 72.9 | 97.1 | 85.40 | 27.38 | 2632.31 | 205.56 |
| GRC_XAN_AD_004 | JS6737 | This study | DI | 75.5 | 74.5 | 98.7 | 72.3 | 95.8 | 63.12 | 14.07 | 19711.20 | 2260.17 |
| GRC_XAN_AD_005 | JS6738 | This study | DI | 85.1 | 84.8 | 99.6 | 84.6 | 99.4 | 100.96 | 22.11 | 1822.21 | 52.37 |
| GRC_XAN_AD_006 | JS6739 | This study | DI | 79.8 | 79.4 | 99.5 | 77.4 | 97.0 | 70.42 | 18.73 | 27450.40 | 1275.94 |
| GRC_XAN_AD_007 | JS6740 | This study | DI | 85.8 | 85.4 | 99.5 | 85.2 | 99.3 | 102.46 | 23.93 | 2439.53 | 31.05 |
| GRC_XAN_AD_008 | JS6741 | This study | DI | 90.2 | 89.8 | 99.6 | 89.6 | 99.3 | 106.47 | 24.73 | 811.21 | 14.02 |
| GRC_XAN_AD_009 | JS6742 | This study | DI | 79.6 | 76.4 | 96.0 | 75.6 | 95.0 | 8.77 | 3.00 | 29337.00 | 3478.76 |
| GRC_XAN_AD_010 | JS6743 | This study | DI | 86.1 | 85.8 | 99.7 | 85.6 | 99.4 | 102.50 | 23.68 | 27.59 | 3.82 |
| GRC_XAN_AD_011 | JS6744 | This study | DI | 80.5 | 80.2 | 99.6 | 80.1 | 99.5 | 96.85 | 23.19 | 23.29 | 1.41 |
| GRC_XAN_AD_012 | JS6745 | This study | DI | 72.3 | 71.9 | 99.4 | 71.8 | 99.3 | 85.20 | 23.45 | 136.61 | 4.07 |
| ITA_NEA_AD_001 | JS6748 | This study | DI | 70 | 69.7 | 99.6 | 69.6 | 99.4 | 55.24 | 15.64 | 26966.10 | 1738.75 |
| ITA_NEA_AD_002 | JS6749 | This study | DI | 66.8 | 66.3 | 99.3 | 66.2 | 99.1 | 74.31 | 25.05 | 9924.74 | 109.39 |
| ITA_NEA_AD_003 | JS6750 | This study | DI | 70.3 | 69.8 | 99.3 | 69.6 | 99.0 | 72.55 | 24.70 | 17578.30 | 250.91 |
| ITA_NEA_AD_004 | JS6751 | This study | Outgroup | 63.9 | 4 | 6.3 | 2.4 | 3.8 | 0.02 | 0.05 | 581.83 | 3.24 |
| ITA_NEA_AD_005 | JS6752 | This study | Outgroup | 32.3 | 3.4 | 10.5 | 1.9 | 5.9 | 0.04 | 0.05 | 511.96 | 1.64 |
| ITA_PAV_AD_001 | NA | Public data | DI | 184.6 | 183.7 | 99.5 | 183 | 99.1 | 10.23 | 2.49 | 208.39 | 14.41 |
| MYS_SEL_AD_001 | JS6763 | This study | DI | 68.1 | 52.3 | 76.8 | 50.3 | 73.9 | 15.72 | 4.59 | 1300.92 | 0.69 |
| MYS_SEL_AD_001_R | JS6764 | This study | DI (REP) | 108.5 | 0.5 | 0.5 | 0.3 | 0.3 | 0.10 | 0.03 | 74.32 | 0.27 |
| PAN_BOC_AD_001 | JS6664 | This study | DI | 77.7 | 77.4 | 99.6 | 77.2 | 99.4 | 86.43 | 21.25 | 3615.37 | 1.14 |
| PAN_BOC_AD_002 | JS6665 | This study | DI | 72.3 | 72 | 99.6 | 62.9 | 87.0 | 51.75 | 17.49 | 24364.90 | 1490.62 |
| PAN_BOC_AD_003 | JS6666 | This study | DI | 71.5 | 71.3 | 99.7 | 71.2 | 99.6 | 81.77 | 22.63 | 3131.66 | 1.24 |
| PAN_BOC_AD_004 | JS6667 | This study | DI | 73.2 | 72.9 | 99.6 | 72.4 | 98.9 | 78.51 | 23.18 | 13772.50 | 248.06 |
| PAN_BOC_AD_005 | JS6668 | This study | DI | 133.4 | 132.8 | 99.6 | 129.5 | 97.1 | 125.24 | 38.68 | 60071.10 | 975.29 |
| PAN_PUE_AD_001 | JS6655 | This study | DI | 71 | 70.7 | 99.6 | 66.2 | 93.2 | 74.81 | 22.59 | 11288.30 | 249.00 |
| PAN_PUE_AD_002 | JS6656 | This study | DI | 75.1 | 74.8 | 99.6 | 74.7 | 99.5 | 90.31 | 22.68 | 2795.07 | 53.89 |
| PAN_PUE_AD_003 | JS6657 | This study | DI | 75.1 | 74.9 | 99.7 | 74.7 | 99.5 | 85.30 | 22.96 | 2336.28 | 51.46 |
| PAN_PUE_AD_004 | JS6658 | This study | DI | 71.2 | 71 | 99.7 | 70.8 | 99.4 | 74.10 | 25.62 | 21454.80 | 422.38 |
| PAN_PUE_AD_004_R | JS6699 | This study | DI (REP) | 99.9 | 99.5 | 99.6 | 97.7 | 97.8 | 104.03 | 36.35 | 4923.96 | 52.06 |
| PAN_PUE_AD_005 | JS6659 | This study | DI | 75.4 | 75.1 | 99.6 | 74.9 | 99.3 | 85.61 | 21.82 | 2290.60 | 52.14 |
| PAN_PUE_AD_006 | JS6660 | This study | DI | 72.8 | 72.5 | 99.6 | 72.4 | 99.5 | 82.19 | 19.37 | 5184.77 | 113.67 |
| PAN_SLO_AD_001 | JS6652 | This study | DI | 75.2 | 74.9 | 99.6 | 74.4 | 98.9 | 79.91 | 27.04 | 20772.80 | 223.07 |
| PAN_SLO_AD_002 | JS6653 | This study | DI | 67.5 | 67.3 | 99.7 | 67.1 | 99.4 | 79.13 | 19.84 | 4577.69 | 89.79 |
| PAN_SLO_AD_003 | JS6654 | This study | DI | 67.8 | 67.4 | 99.4 | 67.2 | 99.1 | 50.42 | 13.84 | 38884.00 | 2216.41 |
| ROU_BUC_AD_001 | JS6671 | This study | DI | 75 | 74.8 | 99.7 | 74.7 | 99.6 | 40.79 | 8.69 | 19945.80 | 4333.07 |
| ROU_COM_AD_001 | JS6675 | This study | DI | 63.4 | 62.7 | 98.9 | 62.5 | 98.6 | 65.06 | 23.97 | 329.45 | 0.67 |
| ROU_GIU_AD_001 | JS6678 | This study | DI | 72.5 | 72.1 | 99.4 | 71.8 | 99.0 | 74.33 | 20.98 | 34666.00 | 71.97 |
| THA_BKK_AD_001 | JS6604 | This study | DI | 78.3 | 78.1 | 99.7 | 77.9 | 99.5 | 90.81 | 29.01 | 17158.90 | 42.36 |
| THA_BKK_AD_002 | JS6605 | This study | DI | 85.7 | 85.5 | 99.8 | 85.3 | 99.5 | 101.44 | 24.38 | 7033.69 | 16.75 |
| THA_BKK_AD_003 | JS6648 | This study | DI | 76 | 75.7 | 99.6 | 74.8 | 98.4 | 82.49 | 23.35 | 25511.60 | 56.19 |
| THA_BKK_AD_004 | JS6649 | This study | DI | 69.3 | 69.1 | 99.7 | 69 | 99.6 | 83.03 | 18.66 | 4130.25 | 4.95 |
| THA_BKK_AD_005 | JS6650 | This study | DI | 80.6 | 80.3 | 99.6 | 80.1 | 99.4 | 93.30 | 26.10 | 7594.70 | 30.82 |
| THA_BKK_AD_006 | JS6651 | This study | DI | 70.8 | 70.6 | 99.7 | 70.4 | 99.4 | 81.78 | 17.01 | 8372.34 | 67.04 |
| USA_FLO_AD_001 | JS6718 | This study | DI | 56.1 | 54.7 | 97.5 | 54.5 | 97.1 | 31.75 | 10.39 | 39173.80 | 2425.45 |
| USA_FLO_AD_002 | JS6719 | This study | DI | 80.5 | 80.2 | 99.6 | 80.1 | 99.5 | 44.48 | 10.95 | 22199.50 | 4823.06 |
| USA_FLO_AD_003 | JS6720 | This study | DI | 102.7 | 102.4 | 99.7 | 100.9 | 98.2 | 100.76 | 25.34 | 37048.40 | 1054.54 |
| USA_FLO_AD_004 | JS6721 | This study | DI | 94 | 93.6 | 99.6 | 93.4 | 99.4 | 109.93 | 25.65 | 1907.00 | 111.26 |
| USA_FLO_AD_005 | JS6722 | This study | DI | 88.7 | 88.4 | 99.7 | 88.2 | 99.4 | 100.88 | 27.82 | 10672.60 | 228.74 |
| USA_FLO_AD_006 | JS6723 | This study | DI | 89.7 | 89.3 | 99.6 | 89.1 | 99.3 | 108.12 | 24.52 | 2820.28 | 99.11 |
| USA_FLO_AD_007 | JS6724 | This study | DI | 74.4 | 74.2 | 99.7 | 74.1 | 99.6 | 24.16 | 6.25 | 23313.10 | 5974.11 |
| USA_FLO_AD_008 | JS6725 | This study | DI | 93.3 | 93 | 99.7 | 92.7 | 99.4 | 113.31 | 24.96 | 1093.87 | 31.24 |
| USA_FLO_AD_009 | JS6726 | This study | DI | 94.6 | 94.2 | 99.6 | 94 | 99.4 | 110.14 | 23.56 | 2540.97 | 73.19 |
| USA_FLO_AD_010 | JS6727 | This study | DI | 97.8 | 97.4 | 99.6 | 97.2 | 99.4 | 113.00 | 26.81 | 8725.98 | 77.97 |
| USA_FLO_AD_011 | JS6728 | This study | DI | 107.6 | 107.2 | 99.6 | 107 | 99.4 | 122.66 | 28.74 | 7542.33 | 248.45 |
| USA_FLO_AD_012 | JS6729 | This study | DI | 96.5 | 96.2 | 99.7 | 95.9 | 99.4 | 102.02 | 25.13 | 5853.22 | 1259.64 |
| USA_FLO_AD_013 | JS6730 | This study | DI | 83.9 | 83.6 | 99.6 | 83.4 | 99.4 | 65.10 | 16.07 | 35588.30 | 2427.30 |
| USA_FLO_AD_014 | JS6731 | This study | DI | 94.5 | 94.2 | 99.7 | 94 | 99.5 | 56.54 | 13.37 | 20280.20 | 5127.06 |
| USA_FLO_AD_015 | JS6732 | This study | DI | 80.4 | 80 | 99.5 | 79.8 | 99.3 | 71.54 | 17.20 | 51591.10 | 715.34 |
| USA_FLO_AD_016 | JS6733 | This study | DI | 77.4 | 77.1 | 99.6 | 76.9 | 99.4 | 92.13 | 24.13 | 2088.03 | 100.71 |
| USA_GEO_AD_001 | NA | Public data | DI | 43.8 | 43 | 98.2 | 42.3 | 96.6 | 26.07 | 7.48 | 6553.15 | 818.36 |
| USA_ILL_AD_001 | JS6597 | This study | DI | 80.9 | 80.7 | 99.8 | 80.6 | 99.6 | 53.45 | 21.90 | 16034.20 | 3774.34 |
| USA_ILL_AD_002 | JS6598 | This study | DI | 91.5 | 91.4 | 99.9 | 91.3 | 99.8 | 30.23 | 7.53 | 26539.90 | 6756.01 |
| USA_LOU_AD_001 | JS6716 | This study | DI | 88.1 | 87.8 | 99.7 | 87.5 | 99.3 | 53.52 | 14.12 | 26351.60 | 4652.53 |
| USA_LOU_AD_002 | JS6717 | This study | DI | 91.5 | 91.1 | 99.6 | 90.9 | 99.3 | 101.77 | 31.93 | 13554.40 | 400.59 |
| USA_MIS_AD_001 | JS6599 | This study | DI | 71.7 | 71.5 | 99.7 | 71.4 | 99.6 | 77.80 | 23.58 | 21256.70 | 416.05 |
| USA_MIS_AD_002 | JS6600 | This study | DI | 106 | 105.7 | 99.7 | 105.6 | 99.6 | 116.98 | 32.89 | 1757.16 | 93.25 |
| USA_TEX_AD_001 | JS6701 | This study | DI | 83.3 | 81.4 | 97.7 | 80.5 | 96.6 | 27.47 | 8.63 | 23184.40 | 5242.61 |
| USA_TEX_AD_002 | JS6702 | This study | DI | 99.2 | 98.9 | 99.7 | 98.1 | 98.9 | 44.81 | 13.27 | 24445.80 | 6107.68 |
| USA_TEX_AD_003 | JS6703 | This study | DI | 79.8 | 79.5 | 99.6 | 79.3 | 99.4 | 94.95 | 22.23 | 4217.91 | 68.45 |
| USA_TEX_AD_004 | JS6704 | This study | DI | 94.3 | 94 | 99.7 | 93.8 | 99.5 | 110.35 | 24.91 | 2851.01 | 70.20 |
| USA_TEX_AD_005 | JS6705 | This study | DI | 97.2 | 96.9 | 99.7 | 96.7 | 99.5 | 110.20 | 24.10 | 2679.54 | 28.78 |
| USA_TEX_AD_006 | JS6706 | This study | DI | 89 | 88.7 | 99.7 | 88.3 | 99.2 | 95.80 | 30.02 | 25003.00 | 89.92 |
| USA_TEX_AD_007 | JS6707 | This study | DI | 97.1 | 96.7 | 99.6 | 96.4 | 99.3 | 112.26 | 26.58 | 2258.08 | 7.63 |
| USA_TEX_AD_008 | JS6708 | This study | DI | 103.2 | 102.8 | 99.6 | 102.5 | 99.3 | 116.18 | 34.72 | 9606.80 | 29.82 |
| USA_TEX_AD_009 | JS6709 | This study | DI | 100 | 99.6 | 99.6 | 99.4 | 99.4 | 112.67 | 28.02 | 2866.40 | 14.18 |
| USA_TEX_AD_010 | JS6710 | This study | DI | 100.4 | 100.1 | 99.7 | 97.9 | 97.5 | 105.34 | 31.17 | 19681.40 | 220.49 |
| USA_TEX_AD_011 | JS6711 | This study | DI | 97.9 | 96.3 | 98.4 | 95.5 | 97.5 | 98.78 | 28.72 | 32060.10 | 522.07 |
| USA_TEX_AD_012 | JS6712 | This study | DI | 68.5 | 67.9 | 99.1 | 67.6 | 98.7 | 70.78 | 17.07 | 33217.60 | 476.19 |
| USA_TEX_AD_013 | JS6713 | This study | DI | 90.4 | 90.1 | 99.7 | 89.7 | 99.2 | 49.91 | 14.87 | 35760.60 | 4540.47 |
| USA_TEX_AD_014 | JS6714 | This study | DI | 87.5 | 87.2 | 99.7 | 86.7 | 99.1 | 100.79 | 30.35 | 24462.40 | 163.93 |
| USA_TEX_AD_015 | JS6715 | This study | DI | 83.1 | 82.7 | 99.5 | 82.4 | 99.2 | 86.74 | 27.39 | 27042.40 | 466.04 |
| USA_WIS_AD_001 | JS6753 | This study | Outgroup | 83.2 | 22.1 | 26.6 | 12.9 | 15.5 | 6.74 | 4.92 | 2864.30 | 90.68 |
| VNM_HAI_AD_001 | JS6747 | This study | Outgroup | 67.7 | 6.6 | 9.7 | 5.1 | 7.5 | 0.07 | 0.04 | 180.72 | 15.40 |
| VNM_HAN_AD_001 | JS6746 | This study | Outgroup | 88.5 | 34.4 | 38.9 | 26.3 | 29.7 | 0.39 | 1.41 | 9394.66 | 1527.76 |
| | | | | | | | | **Average (DI only)** | **75.47** | **20.38** | **13119.78** | **1047.09** |
