## Supplementary Data 2 for "Population genomics reveals an ancient origin of heartworms in canids"

**Supplementary Data 2: Sample metadata for global cohort of adult *Dirofilaria immitis* with outgroups.**

Abbreviations: DI =*Dirofilaria immitis*; REP = Replicate sample

| Sample ID | Original ID | Parasite ID | Accession | Source | Species | Sample type | Country | Region | Host | Parasite stage | Collection date |
|---|---|---|---|---|---|---|---|---|---|---|---|
| AUS_BNE_AD_001 | JS6342 | P4/22-C1 | NA | This study; Swaid Abdullah | *Dirofilaria immitis* | DI | Australia | Brisbane | Dog (*Canis lupus familiaris*) | Adult | 10/5/2017 |
| AUS_BNE_AD_002 | JS6343 | P4/22-E1 | NA | This study; Swaid Abdullah | *Dirofilaria immitis* | DI | Australia | Brisbane | Dog (*Canis lupus familiaris*) | Adult | 08/2021 |
| AUS_BNE_AD_003 | JS6344 | P4/22-F1 | NA | This study; Swaid Abdullah | *Dirofilaria immitis* | DI | Australia | Brisbane | Dog (*Canis lupus familiaris*) | Adult | 9/2/2021 |
| AUS_BNE_AD_003_R | JS6607 | P4/22-F1 | NA | This study; Swaid Abdullah | *Dirofilaria immitis* (REP) | DI (REP) | Australia | Brisbane | Dog (*Canis lupus familiaris*) | Adult | 9/2/2021 |
| AUS_BNE_AD_004 | JS6345 | P4/22-G1 | NA | This study; Swaid Abdullah | *Dirofilaria immitis* | DI | Australia | Brisbane | Dog (*Canis lupus familiaris*) | Adult | 08/2018 |
| AUS_BNE_AD_005 | JS6346 | P4/22-H1 | NA | This study; Swaid Abdullah | *Dirofilaria immitis* | DI | Australia | Brisbane | Fox (*Vulpes vulpes*) | Adult | 3/9/2022 |
| AUS_BNE_AD_006 | JS6347 | P4/22-I1 | NA | This study; Swaid Abdullah | *Dirofilaria immitis* | DI | Australia | Brisbane | Fox (*Vulpes vulpes*) | Adult | 3/9/2022 |
| AUS_BNE_AD_008 | JS6610 | P4/22-F2 | NA | This study; Swaid Abdullah | *Dirofilaria immitis* | DI | Australia | Brisbane | Dog (*Canis lupus familiaris*) | Adult | 9/2/2021 |
| AUS_BNE_AD_009 | JS6611 | P4/22-F3 | NA | This study; Swaid Abdullah | *Dirofilaria immitis* | DI | Australia | Brisbane | Dog (*Canis lupus familiaris*) | Adult | 9/2/2021 |
| AUS_CNS_AD_001 | JS6349 | P3/22-A1 | NA | This study; Carol Esson | *Dirofilaria immitis* | DI | Australia | Cairns | Dog (*Canis lupus familiaris*) | Adult | NA |
| AUS_CNS_AD_002 | JS6350 | P3/22-A2 | NA | This study; Carol Esson | *Dirofilaria immitis* | DI | Australia | Cairns | Dog (*Canis lupus familiaris*) | Adult | NA |
| AUS_LHR_AD_001 | JS6281 | P1/21-E | NA | This study; Duncan Smith | *Dirofilaria immitis* | DI | Australia | Lockhart River/Cooktown | Dog (*Canis lupus familiaris*) | Adult | 11/18/2021 |
| AUS_ROK_AD_001 | JS6606 | P3/23-A1 | NA | This study; Eloise Fenton | *Dirofilaria immitis* | DI | Australia | Rockhampton | Dog (*Canis lupus familiaris*) | Adult | 2023 |
| AUS_SYD_AD_001 | JS5877 | P6/20-E | SRR13154013 | Public data; NCBI | *Dirofilaria immitis* | DI | Australia | Sydney | Dog (*Canis lupus familiaris*) | Adult | 9/11/2020 |
| AUS_SYD_AD_002 | JS5876 | P6/20-D | SRR13154014 | Public data; NCBI | *Dirofilaria immitis* | DI | Australia | Sydney | Dog (*Canis lupus familiaris*) | Adult | 9/11/2020 |
| AUS_SYD_AD_003 | JS5875 | P6/20-C | SRR13154015 | Public data; NCBI | *Dirofilaria immitis* | DI | Australia | Sydney | Dog (*Canis lupus familiaris*) | Adult | 9/11/2020 |
| AUS_SYD_AD_004 | JS5874 | P6/20-B | SRR13154016 | Public data; NCBI | *Dirofilaria immitis* | DI | Australia | Sydney | Dog (*Canis lupus familiaris*) | Adult | 9/11/2020 |
| AUS_SYD_AD_005 | JS5873 | P6/20-A | SRR13154017 | Public data; NCBI | *Dirofilaria immitis* | DI | Australia | Sydney | Dog (*Canis lupus familiaris*) | Adult | 9/11/2020 |
| AUS_SYD_AD_005_R | JS6609 | P6/20-A | NA | This study; University of Sydney | *Dirofilaria immitis* (REP) | DI (REP) | Australia | Sydney | Dog (*Canis lupus familiaris*) | Adult | 9/11/2020 |
| AUS_SYD_AD_006 | JS6277 | P1/21-B1 | NA | This study; University of Sydney | *Dirofilaria immitis* | DI | Australia | Sydney | Dog (*Canis lupus familiaris*) | Adult | NA |
| AUS_SYD_AD_007 | JS6278 | P1/21-B2 | NA | This study; University of Sydney | *Dirofilaria immitis* | DI | Australia | Sydney | Dog (*Canis lupus familiaris*) | Adult | NA |
| AUS_SYD_AD_008 | JS6279 | P1/21-C | NA | This study; University of Sydney | *Dirofilaria immitis* | DI | Australia | Sydney | Dog (*Canis lupus familiaris*) | Adult | NA |
| AUS_SYD_AD_008_R | JS6608 | P1/21-C | NA | This study; University of Sydney | *Dirofilaria immitis* (REP) | DI (REP) | Australia | Sydney | Dog (*Canis lupus familiaris*) | Adult | NA |
| AUS_SYD_AD_009 | JS6280 | P1/21-D | NA | This study; University of Sydney | *Dirofilaria immitis* | DI | Australia | Sydney | Dog (*Canis lupus familiaris*) | Adult | NA |
| AUS_SYD_AD_010 | JS6601 | P1/23-A1 | NA | This study; University of Sydney | *Dirofilaria immitis* | DI | Australia | Sydney | Dog (*Canis lupus familiaris*) | Adult | 4/18/2023 |
| AUS_SYD_AD_011 | JS6602 | P1/23-A2 | NA | This study; University of Sydney | *Dirofilaria immitis* | DI | Australia | Sydney | Dog (*Canis lupus familiaris*) | Adult | 4/18/2023 |
| AUS_SYD_AD_012 | JS6603 | P1/23-A3 | NA | This study; University of Sydney | *Dirofilaria immitis* | DI | Australia | Sydney | Dog (*Canis lupus familiaris*) | Adult | 4/18/2023 |
| AUS_SYD_AD_013 | JS6756 | P5/24-A1 | NA | This study; Ilze Nel | *Dirofilaria immitis* | DI | Australia | Sydney | Dog (*Canis lupus familiaris*) | Adult | 2024 |
| AUS_SYD_AD_014 | JS6757 | P5/24-A2 | NA | This study; Ilze Nel | *Dirofilaria immitis* | DI | Australia | Sydney | Dog (*Canis lupus familiaris*) | Adult | 2024 |
| AUS_SYD_AD_015 | JS6758 | P5/24-A3 | NA | This study; Ilze Nel | *Dirofilaria immitis* | DI | Australia | Sydney | Dog (*Canis lupus familiaris*) | Adult | 2024 |
| AUS_SYD_AD_016 | JS6765 | P6/24-A1 | NA | This study; The University of Sydney | *Dirofilaria immitis* | DI | Australia | Sydney | Dog (*Canis lupus familiaris*) | Adult | 5/10/2024 |
| AUS_SYD_AD_017 | JS6766 | P6/24-A2 | NA | This study; The University of Sydney | *Dirofilaria immitis* | DI | Australia | Sydney | Dog (*Canis lupus familiaris*) | Adult | 5/10/2024 |
| AUS_TVS_AD_001 | JS6351 | P5/22-A1 | NA | This study; Constantin Constantinoiu | *Dirofilaria immitis* | DI | Australia | Townsville | Dog (*Canis lupus familiaris*) | Adult | 01/03/2022 |
| AUS_TVS_AD_002 | JS6352 | P5/22-B1 | NA | This study; Constantin Constantinoiu | *Dirofilaria immitis* | DI | Australia | Townsville | Dog (*Canis lupus familiaris*) | Adult | 01/03/2022 |
| AUS_TVS_AD_003 | JS6353 | P5/22-C1 | NA | This study; Constantin Constantinoiu | *Dirofilaria immitis* | DI | Australia | Townsville | Dog (*Canis lupus familiaris*) | Adult | 03/03/2022 |
| AUS_TVS_AD_004 | JS6354 | P5/22-D1 | NA | This study; Constantin Constantinoiu | *Dirofilaria immitis* | DI | Australia | Townsville | Dog (*Canis lupus familiaris*) | Adult | 21/02/2022 |
| AUS_TVS_AD_005 | JS6355 | P5/22-E1 | NA | This study; Constantin Constantinoiu | *Dirofilaria immitis* | DI | Australia | Townsville | Dog (*Canis lupus familiaris*) | Adult | 4/5/2019 |
| AUS_TVS_AD_006 | JS6356 | P5/22-F1 | NA | This study; Constantin Constantinoiu | *Dirofilaria immitis* | DI | Australia | Townsville | Dog (*Canis lupus familiaris*) | Adult | 12/04/2021 |
| AUS_TVS_AD_007 | JS6357 | P5/22-G1 | NA | This study; Constantin Constantinoiu | *Dirofilaria immitis* | DI | Australia | Townsville | Dog (*Canis lupus familiaris*) | Adult | 04/05/2021 |
| AUS_TVS_AD_008 | JS6358 | P5/22-H1 | NA | This study; Constantin Constantinoiu | *Dirofilaria immitis* | DI | Australia | Townsville | Dog (*Canis lupus familiaris*) | Adult | 28/01/2022 |
| AUS_TVS_AD_009 | JS6359 | P5/22-I1 | NA | This study; Constantin Constantinoiu | *Dirofilaria immitis* | DI | Australia | Townsville | Dog (*Canis lupus familiaris*) | Adult | NA |
| AUS_TVS_AD_010 | JS6360 | P5/22-J1 | NA | This study; Constantin Constantinoiu | *Dirofilaria immitis* | DI | Australia | Townsville | Dog (*Canis lupus familiaris*) | Adult | 23/10/2019 |
| AUS_TVS_AD_011 | JS6368 | P6/22-A1 | NA | This study; Tony Phillis | *Dirofilaria immitis* | DI | Australia | Townsville | Dog (*Canis lupus familiaris*) | Adult | 2022 |
| AUS_TVS_AD_012 | JS6369 | P6/22-A2 | NA | This study; Tony Phillis | *Dirofilaria immitis* | DI | Australia | Townsville | Dog (*Canis lupus familiaris*) | Adult | 2022 |
| AUS_TVS_AD_013 | JS6370 | P6/22-A3 | NA | This study; Tony Phillis | *Dirofilaria immitis* | DI | Australia | Townsville | Dog (*Canis lupus familiaris*) | Adult | 2022 |
| AUS_TVS_AD_014 | JS6612 | P5/22-A2 | NA | This study; Constantin Constantinoiu | *Dirofilaria immitis* | DI | Australia | Townsville | Dog (*Canis lupus familiaris*) | Adult | 01/03/2022 |
| AUS_TVS_AD_015 | JS6613 | P5/22-E2 | NA | This study; Constantin Constantinoiu | *Dirofilaria immitis* | DI | Australia | Townsville | Dog (*Canis lupus familiaris*) | Adult | 4/5/2019 |
| AUS_TVS_AD_016 | JS6614 | P5/22-F2 | NA | This study; Constantin Constantinoiu | *Dirofilaria immitis* | DI | Australia | Townsville | Dog (*Canis lupus familiaris*) | Adult | 12/04/2021 |
| AUS_TVS_AD_017 | JS6615 | P5/22-I2 | NA | This study; Constantin Constantinoiu | *Dirofilaria immitis* | DI | Australia | Townsville | Dog (*Canis lupus familiaris*) | Adult | NA |
| AUS_TVS_AD_018 | JS6616 | P5/22-J2 | NA | This study; Constantin Constantinoiu | *Dirofilaria immitis* | DI | Australia | Townsville | Dog (*Canis lupus familiaris*) | Adult | 23/10/2019 |
| AUS_TVS_AD_019 | JS6617 | P6/22-A4 | NA | This study; Tony Phillis | *Dirofilaria immitis* | DI | Australia | Townsville | Dog (*Canis lupus familiaris*) | Adult | 7/14/1905 |
| CRI_SJO_AD_001 | JS6670 | Duquesa-1 | NA | This study; Alicia Rojas | *Dirofilaria immitis* | DI | Costa Rica | San José | Cat (*Felis catus*) | Adult | NA |
| GRC_XAN_AD_001 | JS6734 | P3/24-A1 | NA | This study; Elias Papadopoulos | *Dirofilaria immitis* | DI | Greece | Thessaloniki and Xanthi | Dog (*Canis lupus familiaris*) | Adult | 2023 |
| GRC_XAN_AD_002 | JS6735 | P3/24-B2 | NA | This study; Elias Papadopoulos | *Dirofilaria immitis* | DI | Greece | Thessaloniki and Xanthi | Dog (*Canis lupus familiaris*) | Adult | 2024 |
| GRC_XAN_AD_003 | JS6736 | P3/24-C1 | NA | This study; Elias Papadopoulos | *Dirofilaria immitis* | DI | Greece | Thessaloniki and Xanthi | Dog (*Canis lupus familiaris*) | Adult | 2023 |
| GRC_XAN_AD_004 | JS6737 | P3/24-D1 | NA | This study; Elias Papadopoulos | *Dirofilaria immitis* | DI | Greece | Thessaloniki and Xanthi | Dog (*Canis lupus familiaris*) | Adult | 2023 |
| GRC_XAN_AD_005 | JS6738 | P3/24-E1 | NA | This study; Elias Papadopoulos | *Dirofilaria immitis* | DI | Greece | Thessaloniki and Xanthi | Dog (*Canis lupus familiaris*) | Adult | 2023 |
| GRC_XAN_AD_006 | JS6739 | P3/24-F1 | NA | This study; Elias Papadopoulos | *Dirofilaria immitis* | DI | Greece | Thessaloniki and Xanthi | Dog (*Canis lupus familiaris*) | Adult | 2023 |
| GRC_XAN_AD_007 | JS6740 | P3/24-G1 | NA | This study; Elias Papadopoulos | *Dirofilaria immitis* | DI | Greece | Thessaloniki and Xanthi | Dog (*Canis lupus familiaris*) | Adult | 2023 |
| GRC_XAN_AD_008 | JS6741 | P3/24-H1 | NA | This study; Elias Papadopoulos | *Dirofilaria immitis* | DI | Greece | Thessaloniki and Xanthi | Dog (*Canis lupus familiaris*) | Adult | 2023 |
| GRC_XAN_AD_009 | JS6742 | P3/24-I1 | NA | This study; Elias Papadopoulos | *Dirofilaria immitis* | DI | Greece | Thessaloniki and Xanthi | Dog (*Canis lupus familiaris*) | Adult | 2023 |
| GRC_XAN_AD_010 | JS6743 | P3/24-J1 | NA | This study; Elias Papadopoulos | *Dirofilaria immitis* | DI | Greece | Thessaloniki and Xanthi | Dog (*Canis lupus familiaris*) | Adult | 2023 |
| GRC_XAN_AD_011 | JS6744 | P3/24-K1 | NA | This study; Elias Papadopoulos | *Dirofilaria immitis* | DI | Greece | Thessaloniki and Xanthi | Dog (*Canis lupus familiaris*) | Adult | 2023 |
| GRC_XAN_AD_012 | JS6745 | P3/24-L1 | NA | This study; Elias Papadopoulos | *Dirofilaria immitis* | DI | Greece | Thessaloniki and Xanthi | Dog (*Canis lupus familiaris*) | Adult | 2023 |
| ITA_NEA_AD_001 | JS6748 | PAR20/2988 | NA | This study; Patrizia Danesi | *Dirofilaria immitis* | DI | Italy | Northeastern Italy | Dog (*Canis lupus familiaris*) | Adult | NA |
| ITA_NEA_AD_002 | JS6749 | PAR19/9727-1 | NA | This study; Patrizia Danesi | *Dirofilaria immitis* | DI | Italy | Northeastern Italy | Fox (*Vulpes vulpes*) | Adult | NA |
| ITA_NEA_AD_003 | JS6750 | PAR19/9727-2 | NA | This study; Patrizia Danesi | *Dirofilaria immitis* | DI | Italy | Northeastern Italy | Fox (*Vulpes vulpes*) | Adult | NA |
| ITA_NEA_AD_004 | JS6751 | 6370/P20 | NA | This study; Patrizia Danesi | *Dirofilaria repens* | Outgroup | Italy | Northeastern Italy | N/A | Adult | NA |
| ITA_NEA_AD_005 | JS6752 | 1400/P22 | NA | This study; Patrizia Danesi | *Dirofilaria repens* | Outgroup | Italy | Northeastern Italy | N/A | Adult | NA |
| ITA_PAV_AD_001 | NA | NA | ERR034941 | Public data; NCBI | *Dirofilaria immitis* | DI | Italy | Pavia | Dog (*Canis lupus familiaris*) | Adult | NA |
| MYS_SEL_AD_001 | JS6763 | DM3-REP | NA | This study; Reuben Sharma | *Dirofilaria immitis* | DI | Malaysia | Selangor | Dog (*Canis lupus familiaris*) | Adult | 2016 |
| MYS_SEL_AD_001_R | JS6764 | DM3 | NA | This study; Reuben Sharma | *Dirofilaria immitis* (REP) | DI (REP) | Malaysia | Selangor | Dog (*Canis lupus familiaris*) | Adult | 2016 |
| PAN_BOC_AD_001 | JS6664 | BC-I-1 | NA | This study; Alicia Rojas | *Dirofilaria immitis* | DI | Panamá | Iroun Boca Chica Óbito Post Tra | Dog (*Canis lupus familiaris*) | Adult | NA |
| PAN_BOC_AD_002 | JS6665 | BC-I-2 | NA | This study; Alicia Rojas | *Dirofilaria immitis* | DI | Panamá | Iroun Boca Chica Óbito Post Tra | Dog (*Canis lupus familiaris*) | Adult | NA |
| PAN_BOC_AD_003 | JS6666 | BC-I-3 | NA | This study; Alicia Rojas | *Dirofilaria immitis* | DI | Panamá | Iroun Boca Chica Óbito Post Tra | Dog (*Canis lupus familiaris*) | Adult | NA |
| PAN_BOC_AD_004 | JS6667 | BC-I-4 | NA | This study; Alicia Rojas | *Dirofilaria immitis* | DI | Panamá | Iroun Boca Chica Óbito Post Tra | Dog (*Canis lupus familiaris*) | Adult | NA |
| PAN_BOC_AD_005 | JS6668 | BC-I-5 | NA | This study; Alicia Rojas | *Dirofilaria immitis* | DI | Panamá | Iroun Boca Chica Óbito Post Tra | Dog (*Canis lupus familiaris*) | Adult | NA |
| PAN_PUE_AD_001 | JS6655 | PA-Ch-Rex-1 | NA | This study; Alicia Rojas | *Dirofilaria immitis* | DI | Panamá | Puerto Armuelles Chiriquí PTY Rex | Dog (*Canis lupus familiaris*) | Adult | NA |
| PAN_PUE_AD_002 | JS6656 | PA-Ch-Rex-2 | NA | This study; Alicia Rojas | *Dirofilaria immitis* | DI | Panamá | Puerto Armuelles Chiriquí PTY Rex | Dog (*Canis lupus familiaris*) | Adult | NA |
| PAN_PUE_AD_003 | JS6657 | PA-Ch-Rex-3 | NA | This study; Alicia Rojas | *Dirofilaria immitis* | DI | Panamá | Puerto Armuelles Chiriquí PTY Rex | Dog (*Canis lupus familiaris*) | Adult | NA |
| PAN_PUE_AD_004 | JS6658 | PA-Ch-Oly-1 | NA | This study; Alicia Rojas | *Dirofilaria immitis* | DI | Panamá | Puerto Armuelles Chiriquí PTY Oly | Dog (*Canis lupus familiaris*) | Adult | NA |
| PAN_PUE_AD_004_R | JS6699 | PA-Ch-Oly-1-REP | NA | This study; Alicia Rojas | *Dirofilaria immitis* (REP) | DI (REP) | Panamá | Puerto Armuelles Chiriquí PTY Oly | Dog (*Canis lupus familiaris*) | Adult | NA |
| PAN_PUE_AD_005 | JS6659 | PA-Ch-Oly-2 | NA | This study; Alicia Rojas | *Dirofilaria immitis* | DI | Panamá | Puerto Armuelles Chiriquí PTY Oly | Dog (*Canis lupus familiaris*) | Adult | NA |
| PAN_PUE_AD_006 | JS6660 | PA-Ch-Oly-3 | NA | This study; Alicia Rojas | *Dirofilaria immitis* | DI | Panamá | Puerto Armuelles Chiriquí PTY Oly | Dog (*Canis lupus familiaris*) | Adult | NA |
| PAN_SLO_AD_001 | JS6652 | EM-Ch-1 | NA | This study; Alicia Rojas | *Dirofilaria immitis* | DI | Panamá | Playa El Manzal San Lorenzo Chiriquí PTY | Dog (*Canis lupus familiaris*) | Adult | NA |
| PAN_SLO_AD_002 | JS6653 | EM-Ch-2 | NA | This study; Alicia Rojas | *Dirofilaria immitis* | DI | Panamá | Playa El Manzal San Lorenzo Chiriquí PTY | Dog (*Canis lupus familiaris*) | Adult | NA |
| PAN_SLO_AD_003 | JS6654 | EM-Ch-3 | NA | This study; Alicia Rojas | *Dirofilaria immitis* | DI | Panamá | Playa El Manzal San Lorenzo Chiriquí PTY | Dog (*Canis lupus familiaris*) | Adult | NA |
| ROU_BUC_AD_001 | JS6671 | Leopard | NA | This study; Georgiana Deak | *Dirofilaria immitis* | DI | Romania | Bucharest | Leopard (*Panthera pardus*) | Adult | 11/2022 |
| ROU_COM_AD_001 | JS6675 | CJ008711 | NA | This study; Georgiana Deak | *Dirofilaria immitis* | DI | Romania | Comana PN | Wildcat (*Felis silvestris*) | Adult | 2021 |
| ROU_GIU_AD_001 | JS6678 | CJ007967 | NA | This study; Georgiana Deak | *Dirofilaria immitis* | DI | Romania | ocolul silvic Giurgiu | Golden jackal (*Canis aureus*) | Adult | 12/2019 |
| THA_BKK_AD_001 | JS6604 | P2/23-A1 | NA | This study; Piyanan Taweethavonsawat | *Dirofilaria immitis* | DI | Thailand | Bangkok | Dog (*Canis lupus familiaris*) | Adult | NA |
| THA_BKK_AD_002 | JS6605 | P2/23-A2 | NA | This study; Piyanan Taweethavonsawat | *Dirofilaria immitis* | DI | Thailand | Bangkok | Dog (*Canis lupus familiaris*) | Adult | NA |
| THA_BKK_AD_003 | JS6648 | P2/23-B1 | NA | This study; Piyanan Taweethavonsawat | *Dirofilaria immitis* | DI | Thailand | Bangkok | Dog (*Canis lupus familiaris*) | Adult | NA |
| THA_BKK_AD_004 | JS6649 | P2/23-B2 | NA | This study; Piyanan Taweethavonsawat | *Dirofilaria immitis* | DI | Thailand | Bangkok | Dog (*Canis lupus familiaris*) | Adult | NA |
| THA_BKK_AD_005 | JS6650 | P2/23-C1 | NA | This study; Piyanan Taweethavonsawat | *Dirofilaria immitis* | DI | Thailand | Bangkok | Dog (*Canis lupus familiaris*) | Adult | NA |
| THA_BKK_AD_006 | JS6651 | P2/23-C2 | NA | This study; Piyanan Taweethavonsawat | *Dirofilaria immitis* | DI | Thailand | Bangkok | Dog (*Canis lupus familiaris*) | Adult | NA |
| USA_FLO_AD_001 | JS6718 | P2/24-E1 | NA | This study; Heather Walden | *Dirofilaria immitis* | DI | USA | Gilchrist County, Florida | Dog (*Canis lupus familiaris*) | Adult | 2018 |
| USA_FLO_AD_002 | JS6719 | P2/24-F1 | NA | This study; Heather Walden | *Dirofilaria immitis* | DI | USA | Gilchrist County, Florida | Dog (*Canis lupus familiaris*) | Adult | 2018 |
| USA_FLO_AD_003 | JS6720 | P2/24-G1 | NA | This study; Heather Walden | *Dirofilaria immitis* | DI | USA | Gilchrist County, Florida | Dog (*Canis lupus familiaris*) | Adult | 2017 |
| USA_FLO_AD_004 | JS6721 | P2/24-H1 | NA | This study; Heather Walden | *Dirofilaria immitis* | DI | USA | Gilchrist County, Florida | Dog (*Canis lupus familiaris*) | Adult | 2017 |
| USA_FLO_AD_005 | JS6722 | P2/24-J1 | NA | This study; Heather Walden | *Dirofilaria immitis* | DI | USA | Gilchrist County, Florida | Dog (*Canis lupus familiaris*) | Adult | 2018 |
| USA_FLO_AD_006 | JS6723 | P2/24-K1 | NA | This study; Heather Walden | *Dirofilaria immitis* | DI | USA | Gilchrist County, Florida | Dog (*Canis lupus familiaris*) | Adult | 2018 |
| USA_FLO_AD_007 | JS6724 | P2/24-O1 | NA | This study; Heather Walden | *Dirofilaria immitis* | DI | USA | Gilchrist County, Florida | Dog (*Canis lupus familiaris*) | Adult | 2019 |
| USA_FLO_AD_008 | JS6725 | P2/24-P1 | NA | This study; Heather Walden | *Dirofilaria immitis* | DI | USA | Gilchrist County, Florida | Dog (*Canis lupus familiaris*) | Adult | 2019 |
| USA_FLO_AD_009 | JS6726 | P2/24-Q1 | NA | This study; Heather Walden | *Dirofilaria immitis* | DI | USA | Gilchrist County, Florida | Dog (*Canis lupus familiaris*) | Adult | 2019 |
| USA_FLO_AD_010 | JS6727 | P2/24-W1 | NA | This study; Heather Walden | *Dirofilaria immitis* | DI | USA | Gilchrist County, Florida | Dog (*Canis lupus familiaris*) | Adult | 2020 |
| USA_FLO_AD_011 | JS6728 | P2/24-X1 | NA | This study; Heather Walden | *Dirofilaria immitis* | DI | USA | Duval County, Florida | Dog (*Canis lupus familiaris*) | Adult | 9/2/2015 |
| USA_FLO_AD_012 | JS6729 | P2/24-Y1 | NA | This study; Heather Walden | *Dirofilaria immitis* | DI | USA | Duval County, Florida | Dog (*Canis lupus familiaris*) | Adult | 10/23/2015 |
| USA_FLO_AD_013 | JS6730 | P2/24-Z1 | NA | This study; Heather Walden | *Dirofilaria immitis* | DI | USA | Duval County, Florida | Dog (*Canis lupus familiaris*) | Adult | 1/23/2015 |
| USA_FLO_AD_014 | JS6731 | P2/24-AA1 | NA | This study; Heather Walden | *Dirofilaria immitis* | DI | USA | Duval County, Florida | Dog (*Canis lupus familiaris*) | Adult | 7/24/2015 |
| USA_FLO_AD_015 | JS6732 | P2/24-AB1 | NA | This study; Heather Walden | *Dirofilaria immitis* | DI | USA | Gilchrist County, Florida | Dog (*Canis lupus familiaris*) | Adult | 2020 |
| USA_FLO_AD_016 | JS6733 | P2/24-AD1 | NA | This study; Heather Walden | *Dirofilaria immitis* | DI | USA | Gilchrist County, Florida | Dog (*Canis lupus familiaris*) | Adult | 2021 |
| USA_GEO_AD_001 | NA | NA | ERR034940 | Public data; NCBI | *Dirofilaria immitis* | DI | USA | Georgia | Dog (*Canis lupus familiaris*) | Adult | NA |
| USA_ILL_AD_001 | JS6597 | JYD-27-A1 | NA | This study; Mike Dzimianski/Andrew Moorhead | *Dirofilaria immitis* | DI | USA | Illinois | Dog (*Canis lupus familiaris*) | Adult | 3/5/2023 |
| USA_ILL_AD_002 | JS6598 | JYD-27-A2 | NA | This study; Mike Dzimianski/Andrew Moorhead | *Dirofilaria immitis* | DI | USA | Illinois | Dog (*Canis lupus familiaris*) | Adult | 3/5/2023 |
| USA_LOU_AD_001 | JS6716 | P1/24-K1 | NA | This study; Guilherme Gomes Verocai | *Dirofilaria immitis* | DI | USA | Winnsboro, Louisiana | Dog (*Canis lupus familiaris*) | Adult | 5/13/2020 |
| USA_LOU_AD_002 | JS6717 | P1/24-K2 | NA | This study; Guilherme Gomes Verocai | *Dirofilaria immitis* | DI | USA | Winnsboro, Louisiana | Dog (*Canis lupus familiaris*) | Adult | 5/13/2020 |
| USA_MIS_AD_001 | JS6599 | MO-A1 | NA | This study; Mike Dzimianski/Andrew Moorhead | *Dirofilaria immitis* | DI | USA | Missouri | Dog (*Canis lupus familiaris*) | Adult | 3/5/2023 |
| USA_MIS_AD_002 | JS6600 | MO-A2 | NA | This study; Mike Dzimianski/Andrew Moorhead | *Dirofilaria immitis* | DI | USA | Missouri | Dog (*Canis lupus familiaris*) | Adult | 3/5/2023 |
| USA_TEX_AD_001 | JS6701 | P1/24-A1 | NA | This study; Guilherme Gomes Verocai | *Dirofilaria immitis* | DI | USA | Alvin, Texas | Ferret (*Mustela putorius furo*) | Adult | 1/6/2021 |
| USA_TEX_AD_002 | JS6702 | P1/24-B1 | NA | This study; Guilherme Gomes Verocai | *Dirofilaria immitis* | DI | USA | College Station, Texas | Dog (*Canis lupus familiaris*) | Adult | 1/13/2018 |
| USA_TEX_AD_003 | JS6703 | P1/24-C1 | NA | This study; Guilherme Gomes Verocai | *Dirofilaria immitis* | DI | USA | DFW, Texas | Dog (*Canis lupus familiaris*) | Adult | 2021 |
| USA_TEX_AD_004 | JS6704 | P1/24-C2 | NA | This study; Guilherme Gomes Verocai | *Dirofilaria immitis* | DI | USA | DFW, Texas | Dog (*Canis lupus familiaris*) | Adult | 2021 |
| USA_TEX_AD_005 | JS6705 | P1/24-D1 | NA | This study; Guilherme Gomes Verocai | *Dirofilaria immitis* | DI | USA | DFW, Texas | Dog (*Canis lupus familiaris*) | Adult | 2021 |
| USA_TEX_AD_006 | JS6706 | P1/24-E1 | NA | This study; Guilherme Gomes Verocai | *Dirofilaria immitis* | DI | USA | Edinburg, Texas | Dog (*Canis lupus familiaris*) | Adult | 1/13/2021 |
| USA_TEX_AD_007 | JS6707 | P1/24-E2 | NA | This study; Guilherme Gomes Verocai | *Dirofilaria immitis* | DI | USA | Edinburg, Texas | Dog (*Canis lupus familiaris*) | Adult | 1/13/2021 |
| USA_TEX_AD_008 | JS6708 | P1/24-F1 | NA | This study; Guilherme Gomes Verocai | *Dirofilaria immitis* | DI | USA | Fort Worth, Texas | Dog (*Canis lupus familiaris*) | Adult | 16/03/2022 or 6/07/2021 |
| USA_TEX_AD_009 | JS6709 | P1/24-F2 | NA | This study; Guilherme Gomes Verocai | *Dirofilaria immitis* | DI | USA | Fort Worth, Texas | Dog (*Canis lupus familiaris*) | Adult | 16/03/2022 or 6/07/2021 |
| USA_TEX_AD_010 | JS6710 | P1/24-G1 | NA | This study; Guilherme Gomes Verocai | *Dirofilaria immitis* | DI | USA | Fort Worth, Texas | Dog (*Canis lupus familiaris*) | Adult | 16/03/2022 or 6/07/2021 |
| USA_TEX_AD_011 | JS6711 | P1/24-H1 | NA | This study; Guilherme Gomes Verocai | *Dirofilaria immitis* | DI | USA | Houston, Texas | Cat (*Felis catus*) | Adult | 1/2/2023 |
| USA_TEX_AD_012 | JS6712 | P1/24-H2 | NA | This study; Guilherme Gomes Verocai | *Dirofilaria immitis* | DI | USA | Houston, Texas | Cat (*Felis catus*) | Adult | 1/2/2023 |
| USA_TEX_AD_013 | JS6713 | P1/24-I1 | NA | This study; Guilherme Gomes Verocai | *Dirofilaria immitis* | DI | USA | Hutchins, Texas | Dog (*Canis lupus familiaris*) | Adult | 5/24/2022 |
| USA_TEX_AD_014 | JS6714 | P1/24-I2 | NA | This study; Guilherme Gomes Verocai | *Dirofilaria immitis* | DI | USA | Hutchins, Texas | Dog (*Canis lupus familiaris*) | Adult | 5/24/2022 |
| USA_TEX_AD_015 | JS6715 | P1/24-J1 | NA | This study; Guilherme Gomes Verocai | *Dirofilaria immitis* | DI | USA | Huntsville, Texas | Cat (*Felis catus*) | Adult | 3/30/2022 |
| USA_WIS_AD_001 | JS6753 | Durs1_DNA | NA | This study; Jeremy Foster | *Dirofilaria ursi* | Outgroup | USA | Wisconsin | American black bear (*Ursus americanus*) | Adult | NA |
| VNM_HAI_AD_001 | JS6747 | Sample 3 (UB-QN) | NA | This study; Dung Thi Bui | *Dirofilaria* sp. 'Thailand II' | Outgroup | Vietnam | Thuy Nguyen, Hai Phong | Human (*Homo sapiens*) | Adult | 2023 |
| VNM_HAN_AD_001 | JS6746 | Sample 2 (B4-103) | NA | This study; Dung Thi Bui | *Dirofilaria* sp. 'Thailand II' | Outgroup | Vietnam | Chuong My, Ha Noi | Human (*Homo sapiens*) | Adult | 2018 |
